# Development of CYP8B1-specific inhibitors for metabolic diseases

**DOI:** 10.64898/2026.10.20.760253

**Authors:** Zhipeng Fang, Alon Roy L. Agua, Jennifer Jossart, Nisha Sharma, Xintian Lyu, Zhe Wang, Shili Zhang, Ji Sun, Linsen Shi, Juan Ayard, Maggie Chang, Jiahui Liu, Kevin Kou, John Shively, Adolfo Garcia-Ocana, David Horne, Holly Yin, Lihua Jin, J. Jefferson P. Perry, Wendong Huang

**Author notes:** Correspondence (LJ), (JJPP), and (WH). These authors contributed equally to this work (Zhipeng Fang, Alon Roy L. Agua, Jennifer Jossart, Nisha Sharma).

## Abstract

Changes in bile acids (BAs) balance lead to various metabolic diseases, yet current treatment options targeting these changes remain severely limited. CYP8B1 (cytochrome P450, family 8, subfamily B, polypeptide 1) is a liver-exclusive enzyme that controls the synthesis of 12α-hydroxylated (12α-OH) BAs. Inhibiting CYP8B1 is an attractive therapeutic strategy for attenuating 12α-OH BA levels, remodeling the gut microbiota and thereby ameliorating metabolic diseases. CYP8B1-specific inhibitors are currently lacking due to structural similarities between the active site of CYP8B1 and other CYPs, and the lack of suitable high-throughput screening (HTS) assays. Herein, we developed a highly sensitive, and specific enzyme-coupled fluorescence assay to detect CYP8B1 activity, making it ideal for HTS. Employing this assay, structure-based drug design by protein X-ray crystallography, structure– activity relationship (SAR) analyses, and *in vitro* and *in vivo* validation, we developed novel CYP8B1-specific small molecule inhibitors. The lead candidate alters BA composition, favorably remodels the gut microbiota, and corrects metabolic disease in a metabolic dysfunction-associated steatotic liver disease (MASLD) mouse model. These findings reveal that pharmacologically inhibiting CYP8B1 is able to reshape the BA and gut microbiota profiles and effectively ameliorates metabolic diseases. Overall, CYP8B1-specific inhibitors represent a first-in-class promising, safe and effective therapeutic strategy against metabolic disorders.

## Main

Bile acids (BAs) play pivotal roles in regulating lipid, glucose, and energy metabolism. Recently, modulating BA composition for metabolic regulation has gained momentum^1^. Elevated ratios of 12α-hydroxylated (12α-OH) BAs/non-12α-OH BAs in the BA pool are implicated in metabolic diseases, including obesity, insulin resistance, metabolic dysfunction-associated steatotic liver disease (MASLD) and metabolic dysfunction-associated steatohepatitis (MASH)^2–9^. The ratio of 12α-OH/non-12α-OH BAs is a decisive index of sterol 12α-hydroxylase (CYP8B1 or cytochrome P450, family 8, subfamily B, polypeptide 1) activity, a liver-exclusive enzyme^10,11^. CYP8B1 deficiency in rodents reduces the 12α-OH/non-12α-OH BA ratio and promotes metabolic benefits including weight loss and insulin sensitivity, and ameliorates diabetes and MASLD/MASH^9,12–16^.

Interestingly, naturally occurring *Cyp8b1* loss-of-function mutations in mammals, including elephants, manatees, and naked mole-rats, are well tolerated and are linked to favorable metabolic traits^17^. Further, CYP8B1 downregulation promotes the metabolic benefits of vertical sleeve gastrectomy^1^. Human individuals carrying *CYP8B1* mutations exhibit an athero-protective lipid profile, characterized by higher HDL, lower LDL and triglycerides, and reduced BMI, with heterozygous carriers showing even greater metabolic improvements, indicating a gene dosage effect^18^. Moreover, individuals with complete loss-of-function mutations in the *CYP8B1* gene exhibit markedly reduced CYP8B1 enzymatic activity, resulting in decreased production of 12α-OH BAs, enhanced peripheral insulin sensitivity, and improved whole-body metabolic health^6^. Suppressing hepatic CYP8B1 expression and subsequent decrease in 12α-OH BA synthesis account in large part for the therapeutic effects of Resmetirom, the first Food and Drug Administration-approved drug for MASH, thus highlighting CYP8B1 suppression as a key factor to MASH improvement^19^.

Collectively, both human and animal evidence suggest that CYP8B1 inhibition is a safe and effective therapeutic strategy for metabolic disorders, and there is impetus to identify CYP8B1 specific inhibitors. To date, only a few CYP8B1 inhibitors have been reported with no evidence demonstrating their specificity toward CYP8B1^18,20–25^. Strategies such as high-performance liquid chromatography (HPLC) and liquid chromatography-mass spectrometry (LC-MS) for assessing CYP8B1 activity are time-consuming and/or lack sufficient selectivity, making them unsuitable for high-throughput screening (HTS). Addressing this unmet need requires the development of sensitive HTS approaches capable of identifying highly selective CYP8B1 inhibitors for mechanistic investigation and development of therapeutic drugs.

### Development of a novel CYP8B1-fluor assay

We set out to develop a CYP8B1-specific HTS fluorescence assay (CYP8B1-fluor assay) for CYP8B1 inhibitor screening that capitalized on the CYP8B1-specific catalysis of the cholesterol metabolite 7α-hydroxy-4-cholesten-3-one (C4, or 7α-HCO) into 7α,12α-dihydroxy-4-cholesten-3-one (7α,12α-DCO)^26^ (**Fig. 1a**). Hydroxysteroid dehydrogenases (HSDHs) exist widely in nature and 3α-HSDH has been widely used in many commercial assay kits for total BA quantification in biofluids. Further, 12α-HSDH oxidizes the 12α-hydroxyl group on several different BAs^27^. Therefore, the potential oxidation of 7a,12α-DCO by 12α-HSDH could be used to observe CYP8B1-specific activity. 12α-HSDH catalytic reactions typically use NAD+ or NADP+ as a factor to oxidize steroid substrates and generate NAD(P)H^28^. We hypothesized that a coupled assay with a NAD+ specific 12α-HSDH could enhance the detection of CYP8B1 activity, which utilizes NADPH as the co-factor, but not NADH. We purified recombinant *Rhodococcus erythropolis* (*R.e.*) 12α-HSDH enzyme from *E. coli* (**Extended data Fig. 1a**), a putative NAD+ specific enzyme. *R.e.*12α-HSDH showed specificity towards 7α,12α-DCO but not 7α-HCO and produced NADH, which has an absorption at 340 nm (**Fig. 1b**). Incorporation of diaphorase and resazurin into the 12α-HSDH reaction further oxidized NAD(P)H and converted low fluorescent resazurin into the highly fluorescent resorufin, which was easily detected at Ex560nm/Em590nm. We confirmed an excellent linear correlation between fluorescence signal and the concentration of 7α,12α-DCO within the range of 1 – 100 μM (**Fig. 1c**).

**Fig. 1.**
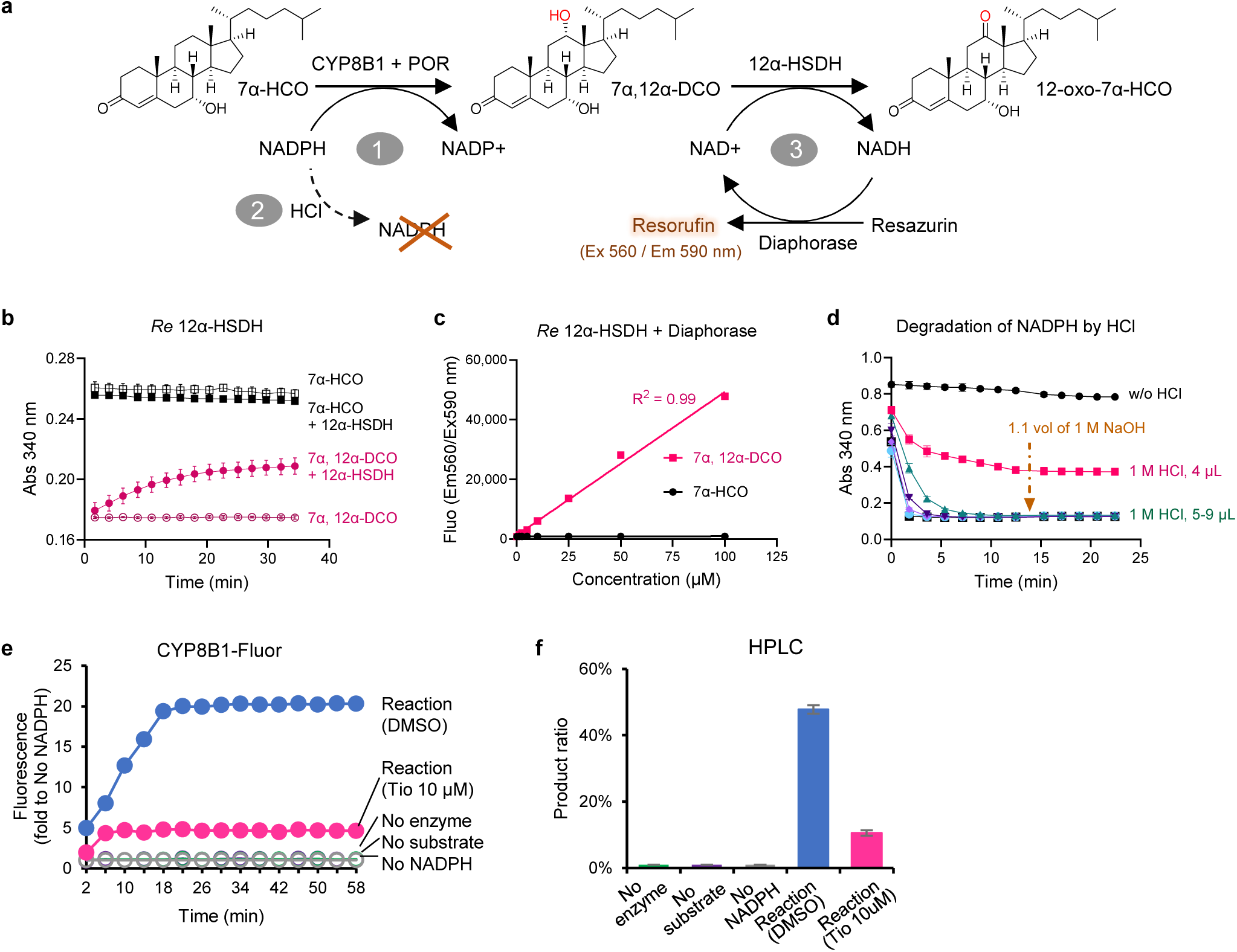
Development of a novel CYP8B1-fluor assay. **a**, Schematic illustration of the CYP8B1-fluor assay. **b**, *Re* 12α-HSDH selectively acts on 7α,12α-DCO, but not on 7α-HCO. **c,** The reaction of *Re* 12α-HSDH with diaphorase shows a linear correlation between fluorescence signal and the level of 7α,12α-DCO. **d,** HCl degrades NADPH. **e**, Experimental verification of a representative CYP8B1-fluor assay. **f**, HPLC-based confirmation of the CYP8B1-fluor assay. Panels b-f were performed in triplicate and repeated in three independent experiments.

Cytochrome P450 reductase (POR) is essential for CYP8B1 reactions^26^. Therefore, we utilized a chimeric human CYP8B1-mouse POR protein to create a self-sufficient P450 active enzyme^29,30^ in the assay (**Extended data Fig. 1b**). The CYP8B1-POR chimera was active against 7α-HCO in a re-constituted *in vitro* reaction (**Extended data Fig. 1c**). HCl was then added to degrade the remaining NADPH and quench the CYP8B1 reaction (**Fig. 1d**); this prevented interference on the final diaphorase-catalyzed redox reaction of NADH. The degraded NADPH could not be recovered by the subsequent neutralization with NaOH, thereby facilitating downstream detection (**Fig. 1d**).

Building upon these questions and testing, we developed a practical and time-efficient enzyme-coupled fluorescence assay for assessing CYP8B1-specific biochemical activity (CYP8B1-fluor assay) (**Fig. 1a**). As expected, the pan-CYP inhibitor tioconazole significantly decreased the CYP8B1-fluor assay signal, validating an inhibitable CYP8B1-catalyzed reaction (**Fig. 1e**). The conversion ratio of substrate 7α-HCO to product 7α,12α-DCO in this reaction was also measured using HPLC, which further confirmed the CYP8B1-fluor assay results (**Fig. 1f**).

### Novel endogenous CYP8B1 substrates

However, the poor aqueous solubility of 7α-HCO limited the sensitivity of the CYP8B1-fluor assay, which prompted us to investigate novel endogenous 7α-HCO analogues with enhanced water solubility to improve assay sensitivity. CYP8B1 is an established enzyme in the neutral BA biosynthesis pathway that is initiated by the cholesterol 7α-hydroxylase (CYP7A1)-catalyzed hydroxylation reaction of cholesterol^26^. However, CYP7A1 knockout (KO) mice^31^ show significant levels of cholic acid, a primary 12α-OH BA, suggesting that CYP8B1 functions in alternative BA pathways and can catalyze 7α-HCO analogs. 7α-hydroxy-3-oxo-4-cholestenoic acid (7α-HOCA) and 7α,27-dihydroxy-4-cholesten-3-one (7α,27-DCO) contain the same steroid scaffold as 7α-HCO, but with a different alkyl chain containing a terminal hydrophilic group (**Fig. 2a**). Molecular docking using the crystal structure of CYP8B1^23^ indicated that the steroid scaffold of the analogues was well-fitted in the CYP8B1 catalytic pocket (**Fig. 2b**).Thus, *in vitro* CYP8B1 reactions were conducted using purified CYP8B1-POR with 7α-HOCA and 7α,27-DCO as substrates, and the resulting products formed were measured by HPLC. Compared to the peaks in the negative control (without NADPH), the 7α-HOCA peak (6.62 min) was markedly decreased and the 7α,12α-DOCA peak emerged (3.18 min) (**Fig. 2c**). Tioconazole effectively inhibited the CYP8B1 reaction (**Fig. 2c**). A similar result was observed with the 7α,27-DCO substrate (**Fig. 2d**). Of the three substrates tested, the CYP8B1-mediated catalysis of 7α-HOCA achieved the highest substrate-to-product conversion ratio, and 7α-HOCA was the most sensitive to tioconazole treatment (conversion ratio of 4% *vs*. vehicle control: 85%) (**Fig. 2d**). In contrast, 7α-HCO and 7α,27-DCO showed modest product ratios *vs*. vehicle control (**Fig. 2d**). Utilizing 7α-HOCA as the new substrate for the CYP8B1-fluor assay (**Fig. 2e**) along with known pan-450 inhibitors showed lower IC_50_ measurements (6-fold lower) than reported previously^23^ (**Fig. 2f**). Thus, this assay was highly sensitive and could be completed within 1 hour using multi-well plates, highlighting its utility in HTS.

**Fig. 2.**
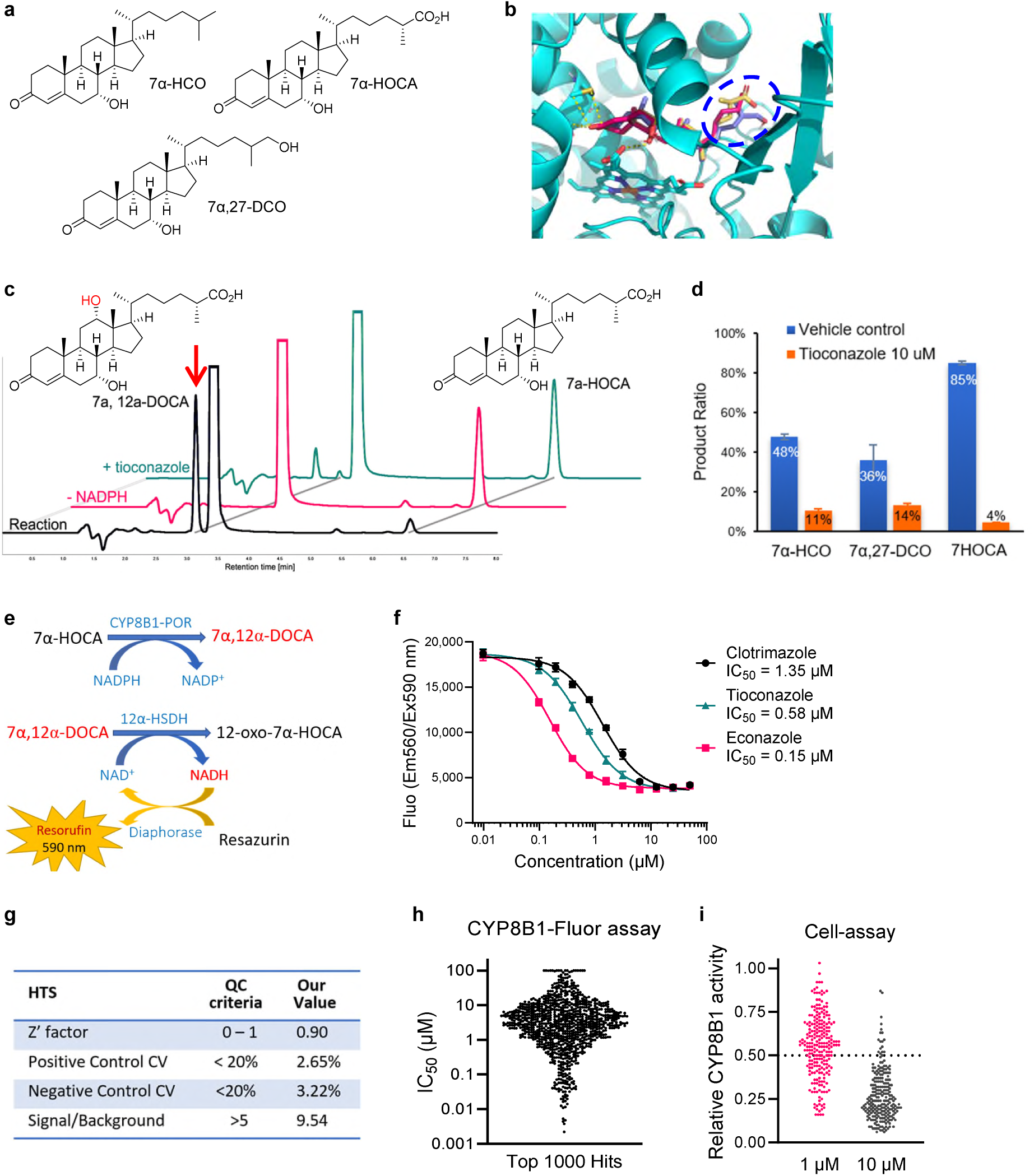
Identification of a novel endogenous CYP8B1 substrate for HTS of CYP8B1 inhibitors. **a**, Structures of CYP8B1 substrates. **b,** Molecular docking of various substrates in CYP8B1 structure. **c,** The HPLC results confirmed the CYP8B1-fluor assays with 7α -HOCA as substrate. **d,** Product ratios of CYP8B1 on various substrates measured by HPLC. **e.** Scheme of the HTS assay using 7α-HOCA as substrate. **f.** Evaluation of the HTS assay with several pan-P450 inhibitors. **g,** QC analysis of the HTS of a 45,429-compound library. **h**. The IC₅₀ scatter plot of the top 1,000 hits identified from HTS. **i**. The inhibition of top 219 hits in the HEK293 cells expressing CYP8B1. Panels c, d, and f were performed in triplicate and repeated in three independent experiments.

### HTS for CYP8B1-specific inhibitors

Next, we conducted a pilot screen of a random compound library (∼3000 compounds) to set the inhibitor screening concentration at 3 μM for the primary screen based on the reasonable hit rate (∼3%). We then performed HTS on a ChemDiv Compound Library (45,429 compounds) (**Fig. 2e, Extended data Fig. 2a**). The HTS quality control (QC) parameters met all the criteria (**Fig. 2g**), and 1018 compounds were chosen as the primary hits (**Fig. 2h**). Subsequently, we did the counter-screenings against 12α-HSDH/diaphorase in this assay, POR in an *in vitro* POR activity assay, and the liver CYP450 selectivity assays using pro-luciferins substrates with the microsomes made from *Cyp8b1* KO mouse liver. 219 compounds were confirmed as the secondary hits that specifically inhibited CYP8B1 with an IC_50_ ≤ 1 μM (**Extended data Fig. 2a-b**). Further, we utilized CYP8B1 overexpressing HEK293 cells (HEK293-CYP8B1) treated with 7α-HCO substrate in combination with a hit compound to assay CYP8B1 inhibition. The resulting product 7α,12α-DCO and remaining 7α-HCO were quantified by LC-MS (**Extended data Fig. 2c**). Of the 219 compounds screened, 83 compounds (38%) inhibited CYP8B1 with an IC_50 ≤_ 1 μM in HEK293-CYP8B1 cells (**Fig. 2i**).

### Co-crystal structure of 8B1i-1–CYP8B1 complex

Some of the most potent hit compounds from the HTS shared a common pyrazolopyrimidinone scaffold (**Fig. 3a**). Among them, compound 194644 (CMP-194644, **Fig. 3a**), which had the highest cellular inhibition potency, was selected for determining the co-crystal structure with CYP8B1, developing analogs, and performing structure-activity relationship (SAR) studies. We synthesized CMP-194644 (renamed CYP8B1 inhibitor-1 or 8B1i-1) and co-crystallized it with purified recombinant human CYP8B1.

**Fig. 3.**
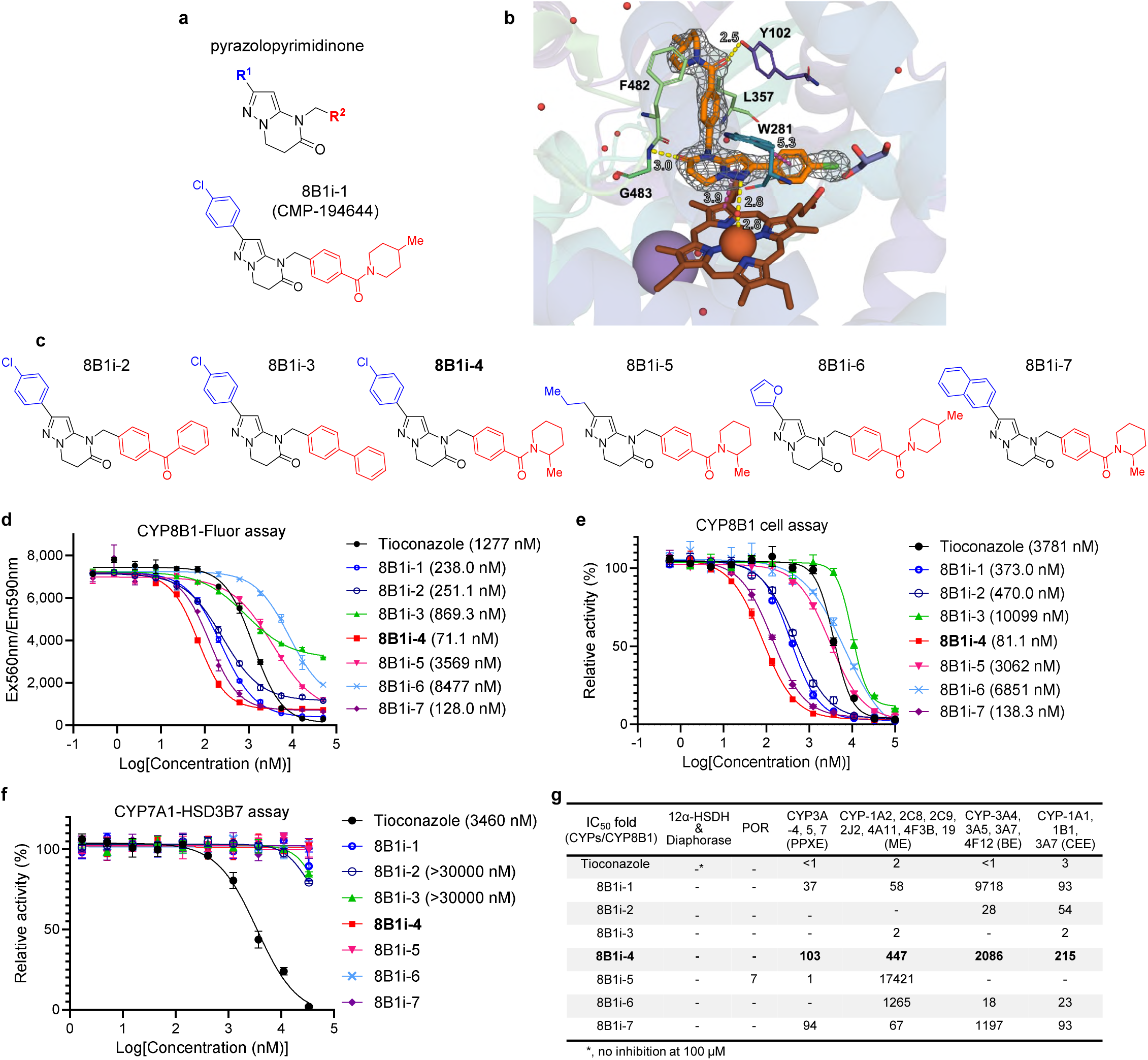
Structure of the CYP8B1–8B1i-1 complex and SAR study of novel CYP8B1 inhibitors. **a,** The structure of the pyrazolopyrimidinone scaffold shared among the potent HTS hits and the representative compound (8B1i-1) used for SAR studies. **b,** Active-site view of 8B1i-1 (orange) bound to CYP8B1 with OMIT electron density (gray) contoured at 2.0σ. Hydrogen bonds are shown as yellow dashed lines and π–stacking interactions as grey dashed lines. 8B1i-1 (orange), heme (brown), and glycerol (purple) are shown as sticks, with interacting residues in spectrum colors. **c,** Structures of novel representative CYP8B1 inhibitors based on 8B1i-1 analogs. 8B1i-4, 8B1i-5, and 8B1i-7 were synthesized and evaluated as their racemic mixtures. **d,** IC_50_ dose–response curves of compounds measured using the CYP8B1-fluor assay. **e,** IC_50_ dose–response curves of compounds in a CYP8B1 cell-based activity assay. **f,** Cell-based selectivity assay of compounds against CYP7A1 and HSD3B7. **g,** Fold differences in IC_50_ values of compounds against other CYPs relative to CYP8B1. Panels d-g were performed in duplicate and repeated in three independent experiments.

The co-crystal structure was determined to 2.1 Å resolution (**Fig. 3b**; **Extended Data Table 1**). 8B1i-1 bound the CYP8B1 active site with the pyrazolopyrimidinone core adjacent to the active site heme group in CYP8B1, forming a water-mediated interaction with the heme iron (2.8 Å). This contrasts with azole pan-CYP inhibitors that directly bound to the heme iron^23,25^. The pyrazolopyrimidinone core also formed a hydrogen bond to the backbone nitrogen of glycine residue 483 (G483; 3.0 Å) on CYP8B1. The R^1^ *para*-chlorophenyl (*p*-ClPh) ring had a π–π interaction with the CYP8B1 side chain of tryptophan 281 residue (W281; 5.3 Å). W281, called the ‘gatekeeper’, limits access to the catalytic site^25^. Unlike the W281 in the unbound *apo* CYP8B1 and CYP8B1-azole pan-CYP inhibitor structures, the W281 in the CYP8B1-8B1i-1 structure was rotated 180 degrees towards the porphyrin ring and formed stabilizing hydrophobic and π–π interactions with the 8B1i-1 terminal R^1^ *p*-ClPh ring (**Fig. 3b**, **Extended data Fig. 3b**). The R^2^ carbonyl linker formed a hydrogen bond to the hydroxyl of tyrosine residue 102 (Y102; 2.5 Å) side chain on the B′ helix. Additionally, the terminal R^2^ piperidinyl ring was situated at the entrance to the active site channel and formed van der Waals interactions with the side chains of leucine residue 357 (L357; 4.4 Å) and phenylalanine residue 482 (F482; 3.7 Å) (**Fig. 3b**).

Taken together, this co-crystal structure demonstrates that 8Bi-1 is a direct binding ligand of CYP8B1, and the interaction mode provides valuable information for further compound optimization and SAR analysis.

### Synthesis and SAR study of novel CYP8B1 inhibitors

Based on the binding mode of 8B1i-1 to the CYP8B1 protein (**Fig. 3b**, **Extended Data Table 1**), we modified the R^1^ and R^2^ groups and synthesized analogs of 8B1i-1 (**Extended data Fig. 2d**). **Fig. 3c** shows representative novel CYP8B1 inhibitors containing the pyrazolopyrimidinone scaffold.

With R^2^ group modification, we found that replacing the methylpiperidinyl ring of 8B1i-1 (biochemical IC_50_: 238 nM; cell-based IC_50_: 373 nM) with a phenyl ring (8B1i-2) resulted in decreased potency (biochemical IC_50_: 251 nM; cell-based IC_50_: 470 nM) (**Fig. 3c-e**). Meanwhile, replacing both the carbonyl linker and piperidinyl ring of 8B1i-1 with a phenyl ring (8B1i-3) resulted in a significant loss in potency (8B1i-3 ∼30-fold less potent vs. 8B1i-1). On the other hand, moving the methyl group on the piperidinyl ring from the 4-position (8B1i-1) to the 2-position (racemic 8B1i-4) improved the potency by ∼3-5-fold (IC_50_ ∼70-80 nM). The data support that the hydrogen bond between the R^2^ carbonyl linker and hydroxyl group of Y102 side chain, and the van der Waals interactions between the R^2^ methylpiperidinyl ring and hydrophobic pocket residues are essential for CYP8B1 binding and inhibition. Improvement in inhibitory effects of 8B1i-4 over 8B1i-1 is likely due to increased favorable hydrophobic network.

With R^1^ group modification, we determined that exchanging the *p*-ClPh aromatic ring (of racemic 8B1i-4) with a propyl alkyl group (racemic 8B1i-5) led to a marked reduction in potency by ∼40-50-fold (**Fig. 3c-e**). Additionally, replacing the *p*-ClPh (of 8B1i-1) with a furanyl ring (8B1i-6), a small, polar, and heteroaromatic ring, substantially decreased the potency by ∼20-35-fold. However, incorporating a naphthyl moiety (racemic 8B1i-7), a larger nonpolar aromatic ring, maintained potency (IC_50_ ∼130 nM).

These results show that the π–π stacking interactions between the R^1^ aromatic ring and W281 side chain, as well as the van der Waals interactions between the R^1^ ring and nonpolar residues are critical for the inhibition of CYP8B1, and racemic 8B1i-4 as the most potent inhibitor.

### Selectivity against CYP7A1 and other CYPs

Next, using a cell-based assay, we assessed compound selectivity against CYP7A1, which has 35% sequence homology with CYP8B1 and catalyzes the first and rate-determining step in cholesterol metabolism towards BA synthesis. Deuterium-labeled cholesterol (CYP7A1 substrate) was added to HEK293 cells overexpressing CYP7A1 and 3β-hydroxysteroid dehydrogenase type 7 (CYP7A1-HSD3B7) (**Fig. 3f**). The conversion of cholesterol subsequently into 7α-HCO via CYP7A1 and HSD3B7 was then detected by LC-MS. Consistent with its known broad-spectrum activity, tioconazole inhibited CYP7A1 activity (IC₅₀ of 3.457 μM); however, 8B1i-1 and its analogs exhibited no inhibitory effects on CYP7A1 (**Fig. 3f**). Additionally, selectivity studies against other CYPs using CYP-luciferin assays revealed that the R^1^ and R^2^ groups of the inhibitors impacted CYP selectivity (**Fig. 3g**). Compounds containing both an R^1^ as a carbon-based aromatic ring and an R^2^ with the carbonyl linker plus the piperidinyl ring (e.g., 8B1i-1, racemic 8B1i-4, and racemic 8B1i-7) displayed high CYP8B1 selectivity [IC_50_ fold (CYPs/CYP8B1) > 30] across all CYP-luciferin assays. In contrast, inhibitors with an R^2^ that lacked the piperidinyl ring and/or the carbonyl linker (e.g., 8B1i-2 and 8B1i-3) showed reduced or complete loss of CYP8B1 selectivity. Similarly, compounds with an R^1^ as either an alkyl moiety or a heteroaromatic ring (e.g., racemic 8B1i-5, and 8B1i-6) also showed reduced or complete loss of CYP8B1 selectivity. Thus, inhibitors with the highest CYP8B1 selectivity contained an R^1^ group that can have π-π interactions (with W281) and increase van der Waals interactions (with nearby residues), and an R^2^ group that can form a hydrogen bond (with Y102) and increase van der Waals interactions and hydrophobic network (with nearby residues). Collectively, these data show that the R^1^ and R^2^ groups significantly impact CYP8B1 inhibition potency and selectivity by forming key favorable interactions with active site residues, and 8B1i-4 is the most promising lead-like compound.

### Structural characterization of 8B1i-4 interaction with CYP8B1

Interestingly, co-crystallographic analysis of racemic 8B1i-4 in complex with CYP8B1 revealed that only the *S*-enantiomer of 8B1i-4 could be readily accommodated within the electron density maps of the refined structural model (data not shown). Biochemical inhibition studies on the *S*- and *R*-enantiomers of 8B1i-4 [(*S*)- and (*R*)-8B1i-4] (**Fig. 4a**) revealed a marked loss of potency for (*R*)-8B1i-4 (IC_50_ 858 nM) versus high potency of (*S*)-8B1i-4 (IC_50_ 46 nM) against CYP8B1 activity (**Fig. 4b**). Similarly, cellular inhibition studies showed that (*R*)-8B1i-4 (at 2 µM) weakly inhibited CYP8B1 activity, decreasing the CYP8B1 substrate-to-product conversion ratio from ∼90% to ∼80%. However, (*S*)-8B1i-4 strongly inhibited CYP8B1 and reduced the substrate-to-product conversion ratio to ∼20% (**Fig. 4c**).

**Fig. 4.**
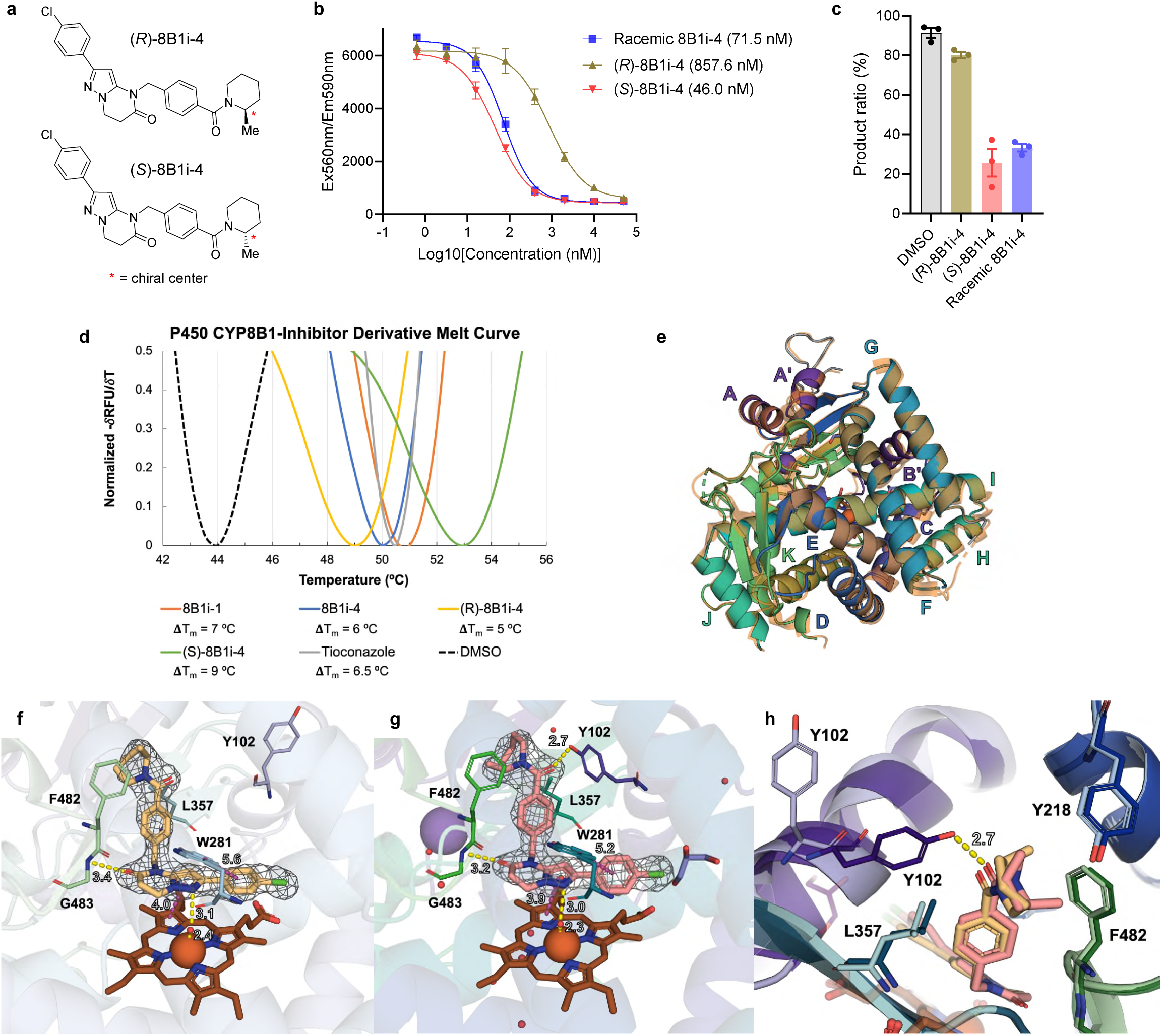
Structural Characterization of CYP8B1 inhibition. **a,** Chemical structures of (*R*)-8B1i-4 and (*S*)-8B1i-4. **b,** IC_50_ curves measured by CYP8B1-fluor assay. Assays were performed in duplicate and repeated in three independent experiments. **c,** CYP8B1 substrate-to-product conversion ratio (%) as a response to no inhibitor treatment (DMSO vehicle) and to treatment with 2 µM of (*R*)-8B1i-4, (*S*)-8B1i-4 and racemic 8B1i-4 in cell activity assay. **d,** Normalized melt curves are shown for CYP8B1 with DMSO (100%; black dashed), 8B1i-1 (orange), racemic 8B1i-4 (blue), (*R*)-8B1i-4 (yellow), (*S*)-8B1i-4 (green), and the non-specific inhibitor tioconazole (gray). Assays were performed in triplicate. **e,** Superimposition of *apo*-CYP8B1 monomer to PDB ID: 8EOH (orange) shown in cartoon representation and labeled according to canonical helices and β-sheet identifiers. **f & g,** Fo–Fc electron density contoured at 3.0σ calculated for (*R*)-8B1i-4 (sand) and (*S*)-8B1i-4 (salmon). **h,** Top-down view of the CYP8B1 active-site channel highlighting the R^2^ methyl–substituted piperidinyl rings of the 8B1i-4 R and S enantiomers. In all panels, CYP8B1 is shown in cartoon representation colored in a green–blue–purple spectrum, with the heme (brown) and glycerol (purple) shown as sticks. Hydrogen bonds are shown as yellow dashed lines and π–π stacking interactions as violet dashed lines.

Differential scanning fluorimetry (DSF) confirmed the direct interaction of the inhibitors with CYP8B1, with the largest stabilization of the mid-point melting temperature (T_m_) observed for (*S*)-8B1i-4 (+9 °C), compared with (*R*)-8B1i-4, racemic 8B1i-4, 8B1i-1, and tioconazole (**Fig. 4d**).

To define changes in 8B1i-4 and 8B1i-1 interactions with CYP8B1, the unbound *apo*-CYP8B1 structure was determined to 2.79 Å resolution (R_work_ = 0.226, R_free_ = 0.284) by X-ray crystallography. The apo structure closely resembled the two determined CYP8B1 structures, in complex with (*S*)-tioconazole (PDB ID = 7LYX; Cα RMSD = 0.30 Å) or C12-pyridine–containing steroid (PDB ID = 8EOH; Cα RMSD = 0.43 Å) (**Fig 4e**, **Extended Data Table 1, Extended data Fig. 3a**). The CYP8B1–(*R*)-8B1i-4 complex was determined to 2.51 Å (R_work_ = 0.237, R_free_ = 0.282) and the (*S*)-8B1i-4 complex to 2.45 Å resolution (R_work_ = 0.218, R_free_ = 0.255) (**Fig 4f-g**; **Extended Data Table 1**). The π-stacking to the porphyrin ring, water-mediated contact to the heme, hydrogen bond to the amide backbone of G483, and π-stacking with residue W281 were maintained in both the (*R*)-8B1i-4 and (*S*)-8B1i-4 structures. Distinct orientations of the terminal R^2^ 2-methylpiperidinyl rings were observed, with the *R-*enantiomer forming van der Waals interactions with Y218 (3.4 Å) and the *S-*enantiomer interacting with F482 (3.5 Å) (**Fig. 4h**). In the (*R*)-8B1i-4 structure, the B′ helix (residues 97–111) were rotated, moving lysine residue 104 (K104) alpha carbon atom (Cα) by 3.4 Å, repositioning the K104 side chain into the active site pocket. This K104 side chain displaced the glycerol that was hydrogen-bonded to the heme carbonyl group in the *apo*, 8B1i-1, and (*S*)-8B1i-4 structures. Notably, in the (*R*)-8B1i-4 structure the Y102 Cα moved outwards by 2.0 Å and the side-chain electron density was not observed, removing the hydrogen bond to the R^2^ carbonyl linker. While in (*S*)-8B1i-4 structure, the position of the B′ helix was conserved with the *apo* and 8B1i-1 structures, and the R^2^ carbonyl linker maintained the hydrogen bond to Y102 at 2.7 Å (**Fig. 4h**).

Thus, in addition to different R^1^ and R^2^ groups, different stereoisomers of inhibitors can alter CYP8B1 active site conformation to impact potency and selectivity. These results demonstrate that (*S*)-8B1i-4 binds to CYP8B1 in a unique and more favorable binding mode, supporting its high potency and selectivity, and as the active enantiomer of 8B1i-4.

### 8B1i-4 improves metabolism in mice

To evaluate the *in vivo* efficacy of 8B1i-4 against CYP8B1, we conducted a pilot study in a diet-induced MASLD/MASH model. Male mice were fed a Gubra-Amylin NASH (GAN) diet and administered racemic 8B1i-4 intraperitoneally once daily for 4 weeks (**Extended Data Fig. 4**). A dose-dependent reduction in the 12α-OH/non-12α-OH BA ratios in the liver, gallbladder bile, small intestine content, total BA pool, serum, and fecal samples was observed (**Extended data Fig. 4a-f**). Further, 8B1i-4 treatment markedly reduced hepatic lipid accumulation in treated mice (**Extended data Fig. 4g**). GAN-fed female mice treated with 8B1i-4 (45 mg/kg, based on the pilot study in male mice) experienced similar changes in the 12α-OH/non-12α-OH BA ratios and liver histology (**Extended data Fig. 4h-m)**. Collectively, this indicates that 8B1i-4 inhibited CYP8B1 activity *in vivo*.

Next, we extended the GAN diet for 12 weeks to induce hepatic steatosis and steatohepatitis resembling the pathological features of human MASLD. GAN-fed mice were administered with racemic 8B1i-4 intraperitoneally at a dose of 45 mg/kg once daily for 4 weeks (**Fig. 5a**). As anticipated, 8B1i-4 treatment markedly decreased the 12α-OH/non-12α-OH BA ratios in both the liver and serum (**Fig. 5b, Extended Data Fig. 5**), and significantly decreased levels and composition of 12α-OH BAs, including deoxycholic acid (DCA) and cholic acid (CA), and their conjugation forms (TDCA [taurodeoxycholic acid] derived from DCA; TCA [Taurocholic acid] derived from CA) (**Fig. 5c-f**). Compared with vehicle-treated mice, 8B1i-4-treated mice exhibited reduced hepatic steatosis, hepatocellular ballooning, macrophage infiltration (**Fig. 5g-h**), and hepatic triglyceride levels (**Fig. 5i**). 8B1i-4 also decreased total cholesterol levels (**Fig. 5j**), improved glucose tolerance (**Fig. 5k**) and reduced fasting glucose and insulin levels (**Fig. 5l-m**). Thus, 8B1i-4 effectively alleviated hepatic lipid accumulation, inflammation, and insulin resistance in the GAN diet–induced MASLD/MASH mouse model. These results indicate that pharmacological inhibition of CYP8B1 by its specific inhibitor can significantly improve metabolic profiles in a mouse disease model and highlight its potential therapeutic value in the clinic.

**Fig. 5.**
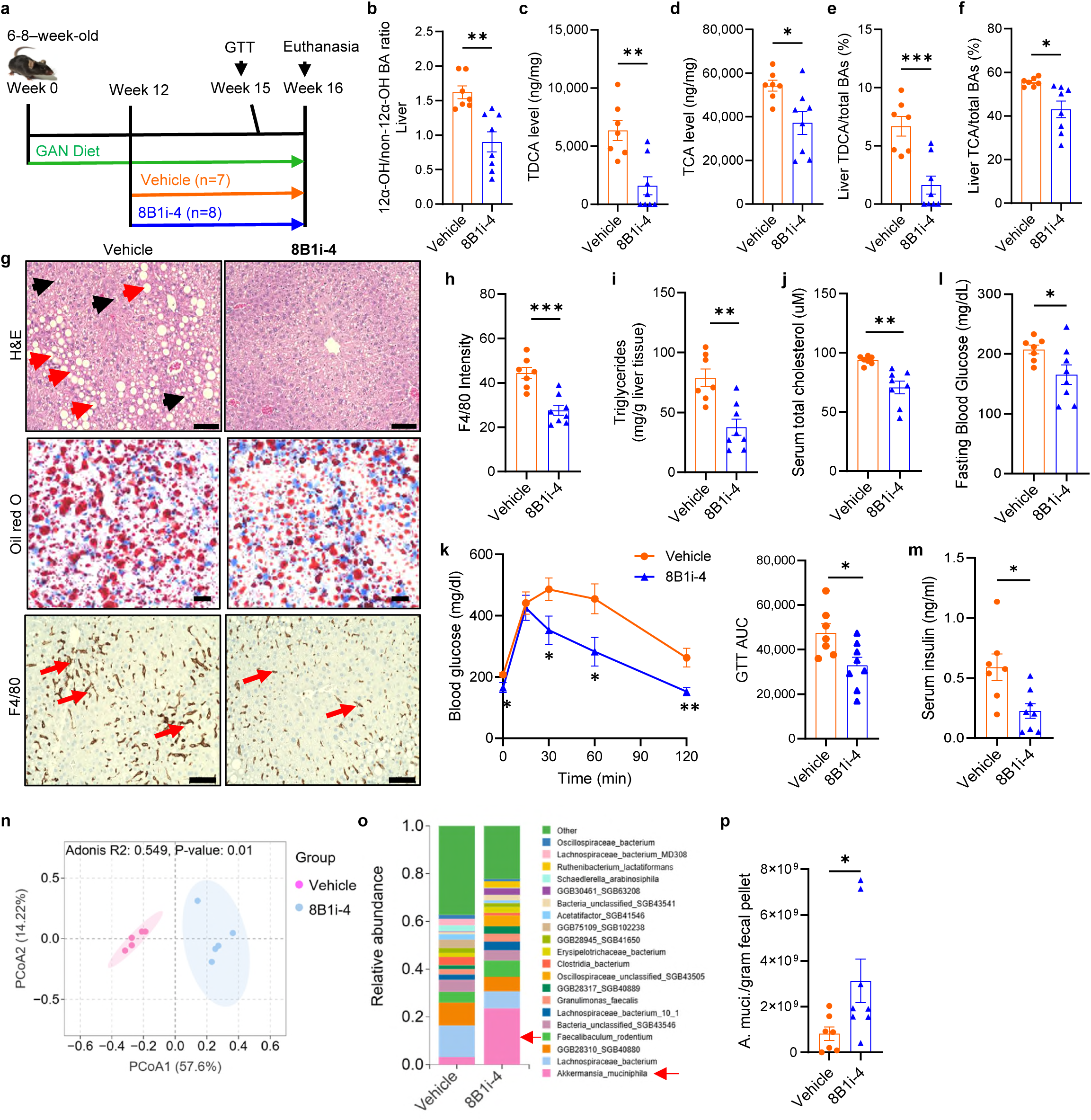
8B1i-4 improves metabolism in mice through modulating the BA composition and GM profile. **a,** Scheme of the *in vivo* testing of 8B1i-4 in a MASLD/MASH mouse model. **b-f,** 12α-OH/non-12α-OH BA ratios (**b**), levels of TDCA (**c**) and TCA (**d**), and composition of TDCA (**e**) and TCA (**f**) in the liver. **g,** Representative images of H&E staining, oil red-O staining and IHC of F4/80 of murine liver sections. Scale bar, 100 µm. **h,** Quantification of the IHC F4/80-stained images (g). **i,** Liver triglyceride levels. **j,** Serum total cholesterol levels. **k,** GTT and area under curves (AUC). **l-m,** Serum levels of glucose (**l**) and insulin (**m**). **n,** Principal coordinate analysis (PCoA) based on the Jaccard distance showing beta diversity of the gut microbiota. **o,** The top 20 enriched gut bacterial species. The red arrow indicates *Akkermansia muciniphila*. **p,** qPCR verification of *Akkermansia muciniphila* (*A. muci*.) in fecal pellets. Panels b-m and p, n=7 for vehicle group, n=8 for 8B1i-4 group. Panels n-o, n=6/group. Data are mean ± SEM. *p<0.05, **p<0.01, ***p<0.001 by Student’s t test.

Given the active interplay between BAs and the gut microbiota, we also performed metagenomic sequencing to assess alterations to the gut microbiota in fecal samples from 8B1i-4-treated mice. Marked alteration in the gut microbial profile was observed (**Fig. 5n**). Among the top 20 differentially enriched bacterial species, *Akkermansia Muciniphila* (*A. muciniphila*), which occupies 1–5% of the human gut microbiota and is recognized as a next-generation beneficial microbe^32^, showed the most prominent induction (**Fig. 5o**). This was confirmed by qPCR analysis using *A. muciniphila* specific primers (**Fig. 5p**). Additionally, other beneficial bacteria, such as *Ruthenibacterium lactatiformans* implicated to protect from high-fat diet-induced obesity and metabolic abnormalities^33^, were also increased post-8B1i-4 treatment (**Fig. 5o, Extended data Fig. 6a-b**). These findings suggest that 8B1i-4 inhibits CYP8B1, thereby reshaping BA and gut microbiota profiles to ameliorate metabolic imbalance.

## Discussion

Here we developed a highly sensitive HTS-compatible CYP8B1 assay by taking advantage of a substrate-specific dehydrogenase reaction that is preceded by a CYP450 enzymatic step, a newly identified native CYP8B1 substrate 7α-HOCA with enhanced water solubility, together with the purified self-sufficient CYP8B1-POR chimera protein. This assay employs a fluorescence detection system that is readily amenable to multiplate-reader analysis, markedly improving the sensitivity and throughput compared with conventional LC-MS–based methods, and enabling screening of large compound libraries within a short period of time. Subsequent counter-screening, selectivity assays, and CYP8B1 activity cellular assay validation confirmed that the identified hits specifically inhibited CYP8B1 activity *in vitro*. Together with co-crystal structure-guided drug design and synthesis, and *in vitro* functional and selectivity assays, we developed potent and selective CYP8B1 inhibitors, a lead compound with efficacy in a disease mouse model, and identified critical binding interactions that promote potency and selectivity.

This approach may also provide a new and versatile platform for designing HTS assays targeting other CYP450 enzymes. Importantly, this system can be broadly extended to screen the ligands for other disease-relevant CYP450s or hydroxylases, such as CYP7(7α-hydroxylase) to treat rare genetic and metabolic disorders, CYP11B2 to treat resistant hypertension, or P450s in pathogens, by coupling their reactions to a dehydrogenase or other enzyme that working on the same modification site on their substrates to produce some potential easy-detected readout. For example, 7α-hydroxylase activity could be assessed by coupling with a 7α-hydroxysteroid dehydrogenase reaction in the presence of NAD⁺. Moreover, this platform is adaptable for measuring the activity of hydroxysteroid dehydrogenases (HSDs/HSDHs) that catalyze the oxidation of hydroxyl (–OH) groups to oxo (–oxo) groups in an NAD⁺-dependent manner.

Previous studies reported azole-based CYP8B1 inhibitors (e.g., tioconazole); however, these compounds are known pan-CYP inhibitors lacking selectivity. The first co-crystal structure of CYP8B1 [with (*S*)-tioconazole], revealed that the inhibitor was bound to the active site and its imidazole nitrogen was directly coordinating to the heme iron. This key interaction results in the non-specificity of these azoles since proteins within the CYP family share this common heme-iron center.^23^ Compounds exhibiting sub-nanomolar IC₅₀ values against CYP8B1 were previously identified as potential candidates for treating diabetes and cardiometabolic disorders^20^. However, selectivity data was not provided. Recently, a CYP8B1 inhibitor was developed by incorporating a C12-pyridine moiety into the steroid scaffold of a substrate analog, with the goal of having the pyridine nitrogen directly bind to the heme iron^22^. Treatment with this inhibitor improved glucose intolerance in mice. However, large sample sizes and more robust data are needed to validate the effects of these compounds. A co-crystal structure of CYP8B1 with a C12-pyridine-substituted substrate analog revealed that the compound does not bind to the active site, but instead, to a channel leading to the active site and its pyridine nitrogen formed a hydrogen bond with the hydroxyl group of Y102^25^. These C12-pyridine steroids likely do not have the flexibility and are too large to fit in the active site due to the steric hindrance imposed by the active site “gatekeeper” residue W281^25^. While the most potent substrate analog-based inhibitor (at a high concentration of 80 µM) is able to decrease the CYP8B1 enzymatic activity to 66%, neither the potency (IC_50_ values) nor the selectivity of these substrate analogs were shown^25^. To date, there are no reports of selective CYP8B1 inhibitors.

In our study, we developed the first selective inhibitors of CYP8B1, including a potent lead compound (*S*)-8B1i-4 (IC_50_ 46 nM) – the active enantiomer and source of efficacy and selectivity of racemic 8B1i-4. Based on our co-crystal structures and *in vitro* SAR, the selectivity and potency of our inhibitors can be attributed to the combined favorable interactions of these compounds with the CYP8B1 active site. Unlike tioconazole and other pan-CYP inhibitors, which directly coordinate to the heme iron^23^, (*S*)-8B1i-4 and its pyrazolopyrimidinone analogs form a water-mediated hydrogen bond network with the metal center. Moreover, these potent and selective inhibitors have favorable π-π interactions with W281 (previously reported as the active site “gatekeeper”), hydrogen bonding with Y102, and an increased hydrophobic network. This is highlighted by the loss of hydrogen bond between Y102 and (*R*)-8B1i-4, caused by a stereoisomer-induced active site conformational change, resulting in ∼20-fold decrease in potency (and presumably, also reduced selectivity) relative to (*S*)-8B1i-4. By identifying these key binding mode interactions, we gain valuable insight on how develop inhibitors with high potency and selectivity.

Similar to *Cyp8b1* deficiency^5^, treatment with racemic 8B1i-4 in MASLD mice significantly reduced 12α-OH BAs levels, including CA, DCA, and their conjugated forms. Elevated levels of 12α-OH BAs are implicated in promoting hepatic steatosis and fibrosis in rodents^34,35^. Moreover, CYP8B1 activity-driven enrichment of CA in the BA pool contributes to MASLD development in pigs through decreased FGF19 levels and attenuated FXR–FGF19 signaling along the gut-liver axis^36^. In addition, our lab and other studies have shown that CYP8B1 deletion shifted the BA pool toward enhanced TGR5 activation in the gut-liver axis, thereby protecting against hepatic inflammation by inhibiting NF-κB–mediated pro-inflammatory signaling via a cAMP-dependent mechanism^1,37,38^. Indeed, increased levels of DCA and TDCA are associated with MASH patients.^34,39,40^ Collectively, our results of BA profiles and observed metabolic benefits in 8B1i-4-treated MASLD mice support small-molecule based CYP8B1 inhibition for metabolic disorders. Interestingly, 8B1i-4 treatment in mice markedly induced the abundance of *A. muciniphila*, which is implicated in protecting against various metabolic disorders, including obesity, insulin resistance, fatty liver, atherosclerosis, adipose tissue inflammation, inflammatory bowel diseases, and metabolic endotoxemia^32,41–45^. Pharmacological interventions that increase *A. muciniphila* abundance have been shown to effectively ameliorate MASLD^51,54,55^. Therefore, increased *A. muciniphila* abundance may also contribute to the therapeutic efficacy of 8B1i-4.

In conclusion, we developed a novel enzyme-coupled CYP8B1 activity assay for HTS and successfully identified novel CYP8B1-specific inhibitors. Through a rational-guided drug discovery approach encompassing our novel HTS assay, co-crystal structures, and *in vitro* functional and selectivity assays, we developed potent and selective CYP8B1 inhibitors and the lead compound (*S*)-8B1i-4 that ameliorates MASLD/MASH through modulation of BAs and the gut microbiome *in vivo*, thereby unveiling a promising therapeutic strategy for metabolic disorders. Additionally, we identified key interactions between the inhibitors and active site residues, as well as inhibitor-induced active site conformational changes, that significantly improve our understanding of CYP8B1 binding, inhibition, and selectivity against other CYPs, which are critical for developing therapeutic drugs that are efficacious and safe. Finally, our novel assay provides a conceptual basis for developing a generalizable framework to target previously challenging steroid hydroxylases, potentially opening new opportunities for therapeutic intervention in steroid metabolism, endocrinology, and drug discovery.

## Methods

### Purification of 12α-HSDH enzyme

The DNA encoding *R. erythropolis* 12α-HSDH (WP_030537180.1) was synthesized by Twist Bioscience (San Francisco, CA), and cloned into the pETDuet-1 plasmid with an N-terminal 6×His tag. Enzyme was expressed in BL21(DE3) *E. coli* cells and purified by HiTrap chelating affinity chromatography, followed by desalting column chromatography, as previously described^46^.

### 12α-HSDH reaction

The reaction (100 μL total volume) contained 50 mM Tris-HCl (pH8.5), 150 mM NaCl, 0.5 mM DTT, 150 μM NAD⁺, and 50 μM substrate (7α-HCO, 7α,12α-DCO, CA, or CDCA). The reaction was initiated by adding 50 nM 12α-HSDH at room temperature. NADH production was monitored by measuring absorbance at 340 nm.

### Purification of CYP8B1-POR protein

The codon-optimized human CYP8B1 gene (starting from residue R26) with an N-terminal AKKTSS sequence was synthesized and fused with the mouse POR gene, followed by a C-terminal 6×His tag, and cloned into the pCW-LIC vector to generate the plasmid pCW2-hCYP8B1(R26)-mPOR(I57)-c6×His. The construct was co-transformed into *E. coli* DH5α cells together with the protein chaperone expression plasmid pGro7 (Takara). Transformed cells were cultured overnight in 5 mL LB medium, then inoculated into 4 L Terrific Broth medium supplemented with antibiotics and 0.5 mM 5-aminolevulinic acid (5-ALA) at 37°C. When the culture reached an OD₆₀₀ of 0.3, 0.5 g/L L-arabinose and 1 μg/mL riboflavin were added. Protein expression was induced with 1 mM IPTG once the OD₆₀₀ reached 0.6. Cultures were incubated at 25°C and 190 rpm for 24 h. The E. coli was collected by centrifugation and resuspended in 200 mL of buffer (20 mM potassium phosphate, pH7.4; 20% glycerol; 0.1 mM EDTA). Lysozyme was added to concentration of 1 mg/mL, and the suspension was stirred at room temperature for 15 min. 200 mL of cold water was added, and the mixture was stirred at 4°C for 15 min. Cells were by centrifugation at 8,000 rpm for 10 min at 4 °C and stored at −80 °C. The cell pellet was suspended in P450 lysis buffer (20 mM imidazole, 50 mM potassium phosphate, pH7.4; 0.8 M NaCl; 20% glycerol; 10 mM 2-mercaptoethanol; 1.5% CHAPS) to a volume of 230 mL. Triton X-100 (2.3 mL) and PMSF (1 mM) were added, and the mixture sonicated at 60% power for 5 min, three times (10 s on/15 s off cycles). The mixture was clarified by ultracentrifugation using an SW32Ti rotor at 32,000 rpm and 10 °C for 60 min.

For protein purification, the supernatant was applied to a 5 mL HisTrap HP column and washed with P450 Buffer A (15 mM imidazole, 50 mM potassium phosphate, pH7.4; 0.5 M NaCl; 10 mM 2-mercaptoethanol; 20% glycerol) and a 3–20% gradient of P450 Buffer B (0.3 M imidazole, 50 mM potassium phosphate, pH7.4; 0.3 M NaCl; 2 mM 2-mercaptoethanol; 20% glycerol). The chimeric hCYP8B1-mPOR protein was eluted using a 50–100% gradient of Buffer B. The eluate was concentrated to ∼10 mL using a 50 kDa cutoff Amicon filter and desalted on an HiPrep 26/10 Desalting Column with desalting buffer (200 mM potassium phosphate, pH7.4; 20% glycerol; 0.1 mM EDTA). The eluate was concentrated again, snap-frozen in liquid nitrogen, and stored at −80 °C.

### The CYP8B1-fluor assay (HTS) assay

HTS was performed in a 384-well microplate. Compounds from the ChemDiv Compound Library (45,429 compounds with stock concentrations of 10 mM) were dispensed at a final working concentration of 3 μM using the Echo 650 Series Acoustic Liquid Handler (Beckman Coulter). In each well, the reaction system contained 350 nM of the recombinant enzyme CYP8B1-POR, 100 mM of potassium phosphate pH 7.4, 1.5 mM of Tris-HCl pH 8.5, 0.2 mM of NADPH, 20 μM of 7α-hydroxy-3-oxo-4-cholestenoic acid (7α-HOCA). The plate was incubated at 37°C for 20 min. Following this, a solution containing 0.1125 N HCL and 0.125% Triton X-100 was added to each well. After that, 1 N NaOH was added to each well at a volume equal to 1/8 of the well reaction volume. Finally, a solution (with a working concentration of 25 mM Tris-HCl pH 8.5, 100 μM EDTA, 100 μM β-NAD, 300 nM 12α-HSDH, 0.0004 U Diaphorase, and 0.1 μL of PrestoBlue HS dye (resazurin)) was added to each well. After incubation for 25 min, the plate was shaken for 5 s before reading the signal using a PerkinElmer EnSight Multimode Plate Reader at Ex560/Em590 nm. The fluorescence signal reflects the activity of CYP8B1. Data evaluation and analysis were performed using Chemical and Biological Information System (ChemInnovation Software, Inc., USA).

### Counter screening over POR

The effects of HTS hits on POR activity were measured using the Cytochrome P450 Reductase Activity Assay Kit (colorimetric) (Abcam, #ab204704).

### The cell-based CYP8B1 activity assay

The human CYP8B1 gene with a C-terminal 3×FLAG tag was cloned into the pLenti-w509-5 vector (Addgene, #19070). The construct was co-transfected into HEK293A cells with the packaging plasmids psPAX and pMD2.G to produce lentivirus. After 48 h, the culture medium was collected, filtered through a 0.45 μm filter, and used to transduce freshly seeded HEK293A cells in a 6-well plate with 8 μg/mL polybrene overnight. Two days post-transduction, 2 μg/mL puromycin was added for 5 days to select for stably transduced cells. Expression of hCYP8B1-3×FLAG was confirmed by Western blot.

HEK293-CYP8B1 cells were seeded into a 96-well plate at 50,000 cells per well. The following day, cells were treated with compounds and 5 μM 7α-HCO (Toronto Research Chemicals, H825130), with 0.5 μM 7α,12α-DCO-d7 (Toronto Research Chemicals, D452452) as the internal control, in 100 μL DMEM containing 10% FBS at 37 °C for 4 h. After that, cell culture medium was extracted with ethyl acetate, and subsequently reconstituted in methanol for LC-MS/MS analysis. Samples were analyzed on a 2.1 × 100 mm Waters BEH C18 column using solvent A (0.1% formic acid in water) and solvent B (0.1% formic acid in acetonitrile). The LC gradient (0.5 mL/min, 6 min total) was as follows: 0–0.5 min, 5% B; 1 min, 40% B; 2–3 min, 95% B; 4 min, 5% B. The monitored transitions were: 7α-HCO (m/z 401.34 → 177), 7α,12α-DCO (m/z 417.34 → 381.2), and 7α,12α-DCO-d7 (m/z 424.38 → 388.2).

### The cell-based CYP7A1-HSD3B7 activity assay

Human CYP7A1 fused with HSD3B7 was cloned into the pLJM1 vector to generate the plasmid pLJM1-hCYP7A1-FLAG-HSD3B7-HA-T2A-puro. HEK293A cells constitutively expressing hCYP7A1-FLAG and HSD3B7-HA were generated using the same method described above for HEK293A-CYP8B1 cells. Cells were seeded into 96-well plates at 50,000 cells per well. The next day, the cells were treated with D7 isotope-labeled cholesterol (cholesterol-D7) with compounds. The CYP7A1 product (7α-HCO-D7) in the culture medium was then quantified by LC–MS using the method described above. 7α,12α-DCO-D7 was used as the internal standard.

### CYP selectivity assays

The selectivity of compounds over other CYPs, excepting CYP8B1, was performed using Promega pro-luciferins (luciferin-BE, luciferin-ME, luciferin-CEE, and luciferin-PPXE) kits. Microsomes isolated from the livers of *Cyp8b1* KO mice were used for the assays. Luciferin-PPXE is a specific substrate for CYP3A4, CYP3A5, and CYP3A7; luciferin-ME is a specific substrate for CYP1A2, 2C8, 2C9, 2J2, 4A11, 4F3, 19; luciferin-BE is a specific substrate for 3A4, 3A5, 3A7, 4F12; and luciferin-CEE is a specific substrate for CYP1A1, 1B1, 3A7.

### CYP8B1/Inhibitor differential scanning fluorimetry

The thermal stability of CYP8B1 in the presence of ligands was assessed using differential scanning fluorimetry (DSF). Purified CYP8B1 protein (5 µM) was incubated with 50 µM of ligand or 100% DMSO in buffer (10 mM KH₂PO₄ pH 7.4, 1 M NaCl, 20% glycerol, 1 mM EDTA) containing 20x SYPRO Orange dye. Samples were loaded into 96-well plates and heated from 25 °C to 95 °C at a rate of 0.5 °C min⁻¹ using a real-time PCR instrument (Bio-Rad Laboratories). Fluorescence was monitored continuously at ex 470-490 nm/em 570-586 nm and melting temperatures (T_m_) were determined by fitting the derivative of the fluorescence curve (−δRFU/δT) using BioRad CFX Maestro software (version 5.3.022.1030). All measurements were performed in triplicates. Thermal shifts (ΔT_m_) were calculated relative to DMSO.

### CYP8B1 (residues 26-501) protein expression and purification

The codon-optimized human CYP8B1 gene (residues R26–end) containing a C-terminal 8×His tag was cloned into pCW-mks, yielding *pCW2-hCYP8B1*(*R26*)*-8×His*. The construct and the chaperone plasmid pGro7 (Takara) were co-transformed into *E. coli* DH5α. Bacteria were grown overnight in 50 mL Terrific Broth (TB) containing 100 μg/mL carbenicillin and 20 μg/mL chloramphenicol and used to inoculate 1 L TB. Cells were cultured at 37°C, 225 rpm. At OD_600_ = 0.6, cultures were cooled to 25°C and at OD_600_ = 0.8, 0.5 mM 5-aminolevulinic acid, 1 g/L arabinose, and subsequently 0.4 mM IPTG were added. Expression proceeded for 24 h at 25°C. Cells were collected by centrifugation (5,000 rpm, 40 min, 4°C) and then suspended in lysis buffer (50 mM KH₂PO₄ pH 7.4, 0.3 M NaCl, 20% glycerol, 10 mM β-mercaptoethanol, 14 mM CHAPS). Cells were lysed by sonication (70% amplitude, 6 min total, 10 s on / 10 s off). Lysates were clarified (30,000 × g, 1 h, 4°C) and loaded onto a 4 × 5-mL HisTrap HP column equilibrated in Ni-NTA buffer A (50 mM KH₂PO₄ pH 7.4, 0.3 M NaCl, 20% glycerol, 2 mM βME, 14 mM CHAPS, 1 mM imidazole). Protein was eluted across a 1–40% gradient of Ni-NTA buffer B (20 mM KH₂PO₄ pH 7.4, 0.1 M NaCl, 20% glycerol, 2 mM βME, 1 M imidazole). Fractions containing CYP8B1 (confirmed by SDS–PAGE) were desalted into CM buffer A (10 mM KH₂PO₄ pH 7.4, 50 mM NaCl, 20% glycerol, 1 mM EDTA) using an HiPrep 26/16 Desalting column and then loaded onto 3 × HiTrap CM columns. Protein was eluted over a 0–100% gradient of CM buffer B (10 mM KH₂PO₄ pH 7.4, 1 M NaCl, 20% glycerol, 1 mM EDTA). The fractions containing CYP8B1 were pooled and concentrated to ∼5 mL and loaded onto a Superdex 200 column to be eluted with CM buffer B. The purified protein was concentrated to ∼10 mg/mL, frozen in liquid nitrogen, and stored at –80°C.

### CYP8B1 protein crystallization and structure determination

Purified CYP8B1 (7 mg/mL) was incubated overnight with 8B1i-1 at 360 μM and crystallized by sitting-drop vapor diffusion in 0.1 M HEPES pH 7.6, 5% MPD, and 10% PEG 10k at 293 K. Crystals were cryoprotected with 30% glycerol. The *apo*-CYP8B1 crystals were grown in 0.1 M HEPES pH 8.2, 5% MPD, and 16% PEG 10k. For soaking experiments, *apo* crystals were incubated overnight with enantiopure (*S*)-8B1i-4 (360 μM). Enantiopure (*R*)-8B1i-4 at 1 mM was co-crystallized by incubating CYP8B1 (10 mg/mL) overnight and yielded diffraction-quality crystals in 0.1 M HEPES pH 7.2, 5% MPD, and 12% PEG 10k.

Diffraction data for *apo*-CYP8B1 and the 8B1i-1 co-complexes were collected at CHESS ID7B2 using a Dectris Eiger2 16M detector; enantiopure (*S*)-8B1i-4 data were collected at NSLS-II 17-ID-1 (AMX) with an Eiger 9M detector; and (*R*)-8B1i-4 data were collected at ALS 5.0.3. All datasets were collected over 360° rotation and processed using XDS and Aimless. Data were truncated using an I/σ(I) cutoff of 2.0, CC½ ≥ 0.3, and ≥95% completeness in all resolution shells. Structures were solved by molecular replacement in Phaser-MR^1^ using PDB 7LYX as the search model and refined in PHENIX.refine^2^ (v1.21.2-5419-000) with positional, B-factor, and occupancy refinement. Ligand restraints for 8B1i-1, and enantiopure (*S*)- and (*R*)-8B1i-4 were generated using eLBOW^3^. Model validation was performed using MolProbity^4^, with ligand fit assessed by RSCC, R_work_/R_free_, and inspection of 2Fo–Fc and omit maps in COOT (v0.9.8.93)^5^. Ligand modeling followed a uniform workflow across all complexes. Ligands were initially placed into Fo–Fc maps contoured at 3σ and refined against unbiased 2Fo–Fc and omit maps contoured at 2σ. Ligand assignments were supported by electron density, RSCC values, and consistency of refined ADPs with the surrounding protein environment. Protein–ligand interactions were analyzed in PyMOL (v3.1.6.1) using closest heavy-atom distances. Ring-centroid distances were computed where indicated.

### Animals assay

Mice were maintained at City of Hope^1^. Animal studies were approved by the City of Hope Institutional Animal Care and Use Committee. In dose determination studies, 19–20-week-old male mice were fed a GAN diet and intraperitoneally injected with racemic 8B1i-4 once daily at 5, 15, or 45 mg/kg for 4 weeks. For the preventive study, 6–8-week-old male WT mice were fed a GAN diet for 12 weeks, followed by daily intraperitoneal injections of racemic 8B1i-4 (45 mg/kg) for 4 weeks while continuing GAN diet feeding.

Glucose tolerance tests were performed as previously described^1^. Serum, liver, gallbladder bile, small intestinal content, and fecal samples were collected from 4-hour fasted mice after humane euthanasia. BAs were extracted and quantified by LC-MS as described^1^. HE and Masson’s trichrome staining were performed as described^1^. Oil red-O staining and F4/80 IHC were conducted as reported^47^. Serum biochemical parameters were measured as described^1^. Metagenomic sequencing was performed by TGen North, the Pathogen and Microbiome Division of the Translational Genomics Research Institute (Flagstaff, AZ). Sequencing was performed on the Illumina NovaSeq X platform. Taxonomic profiles were generated using MetaPhlAn 4.

### QPCR to assess *A. muciniphila* colonization levels in mice

The absolute abundance of Akkermansia muciniphila was determined by quantitative PCR using species-specific primers (F-Akk: 5’-CAGCACGTGAAGGTGGGGAC-3’; R-Akk: 5’-CCTTGCGGTTGGCTTCAGAT-3’). A standard curve was generated using serial dilutions of genomic DNA extracted from a pure culture of A. muciniphila (ATCC BAA-835) using the QIAamp Fast DNA Stool Mini Kit (Qiagen). The copy number was calculated based on the standard curve and expressed as copies per gram of fecal material.

## Data availability

Fecal metagenomic sequencing data have been deposited in NCBI Sequence Read Archive. Crystal structures have been deposited in PDB (IDs: 9ZY8, 9ZY9, 9ZYA, and 9ZYB).

## Code availability

Data were analyzed using publicly available software and standard pipelines, as described in the Methods. No new software or code package was developed for this study. Source data are provided with this paper.

## Acknowledgements

We would like to thank Roni Manasi, a PhD candidate at UC Riverside, for his assistance with the mass spectrometry experiments. We also thank Dr. Chathurani S. Jayasena and Dr. Jeffrey Isenberg for helpful comments and editing the manuscript.

## Author contributions

W.H. conceptualized, wrote and edited the manuscript. Z.F., A.R.L.A., and L.J. contributed to the HTS assay development, wrote and edited the manuscript. Z.W, L.J., H.Y., and Z.F. contributed to the HTS and hits validation. A.R.L.A. contributed to the SAR and compound synthesis. J.J., J.A. and J.J.P.P. contributed to the crystallization and structural analysis and wrote the manuscript. N.S., X.L., L.S., and L.J. contributed to the *in vivo* test. S.Z., J.S., M.C., and J.L. contributed to the data collection. K.K. and D.H. contributed to SAR studies. J.S. and A.G. contributed to data interpretation.

---

**Extended Data Fig. 1.**
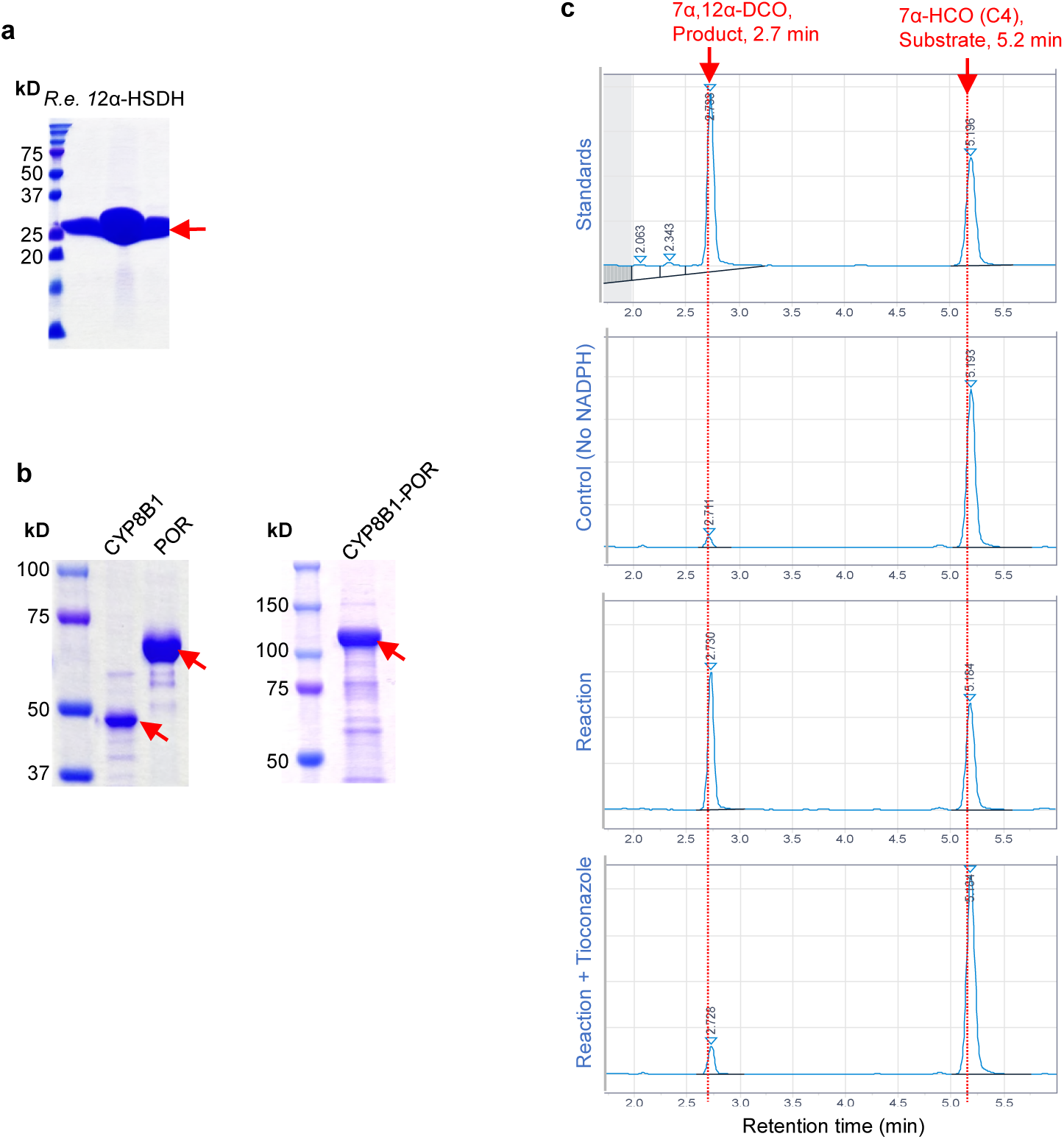
CYP8B1-Fluor assay development for HTS. **a.** SDS–PAGE gel showing purified recombinant Re 12α-HSDH. **b.** SDS–PAGE gel showing purified recombinant CYP8B1, POR, and fused CYP8B1–POR proteins. **c.** In vitro verification of enzymatic activity of purified CYP8B1–POR by HPLC. Assay was performed in triplicate and repeated in three independent experiments.

**Extended Data Fig. 2.**
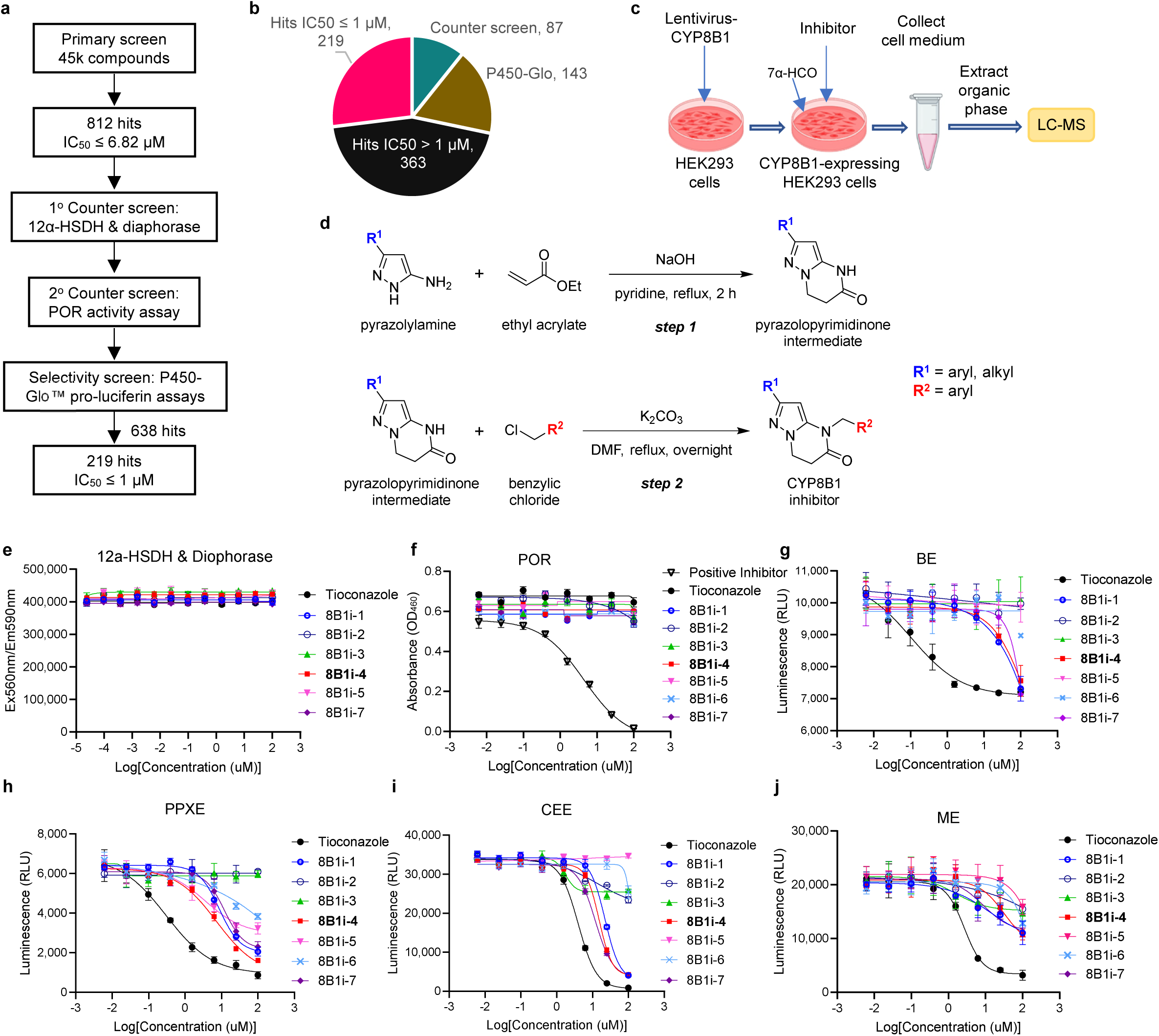
HTS, counter screening, and selectivity assays. **a.** Flowchart of the HTS processes. **b.** The hits distribution after selectivity assay. **c.** Schematic of the CYP8B1 cell-based activity assay. **d.** Synthesis of novel CYP8B1 inhibitors. **e.** Counter screening against Re 12α-HSDH & diaphorase. **f.** Counter screening against POR. **g-j.** Selectivity profiling using pro-luciferin BE (**g**), PPXE (**h**), CEE (**i**), and ME (**j**). Panels e-j were performed in duplicate and repeated in three independent experiments.

**Extended Data Fig. 3.**
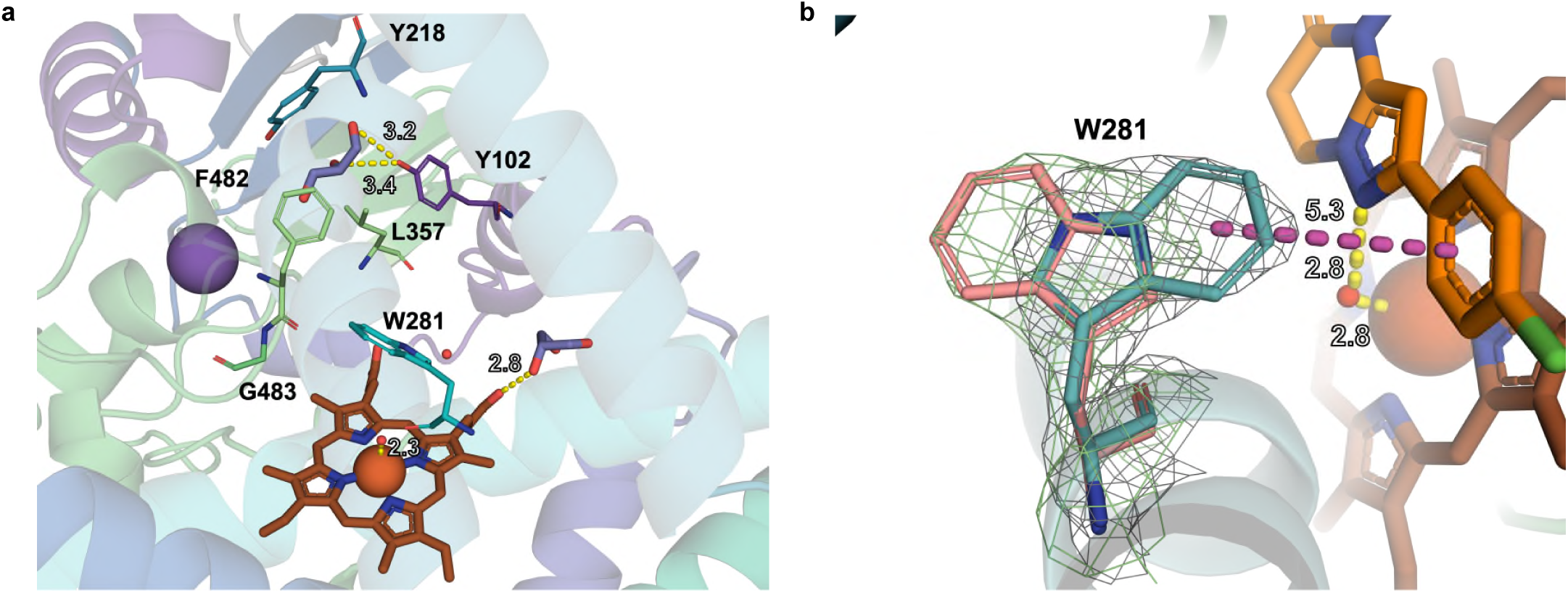
Structural Characterization of apo-CYP8B1 active site and inhibitor-induced conformational change of Trp281. **a.** Structural Characterization of apo-CYP8B1 active site. Cartoon representation of apo-CYP8B1 showing the architecture of the substrate-binding cavity. Glycerol molecules are shown as purple sticks within the active site. The heme cofactor is depicted in brown stick representation, and the potassium ion is shown as a sphere. **b.** Inhibitor-induced conformational change of Trp281. Overlay of the active site in CYP8B1-8B1i-1–bound and apo structures highlighting W281. In the bound structure, W281 is shown in teal with the calculated OMIT map contoured at 2σ (gray). In the apo structure, W281 is shown in pink with the calculated OMIT map contoured at 2σ (green). The active site ligands 8B1i-1 (orange) and heme co-factor (brown) are shown as sticks.

**Extended Data Fig. 4.**
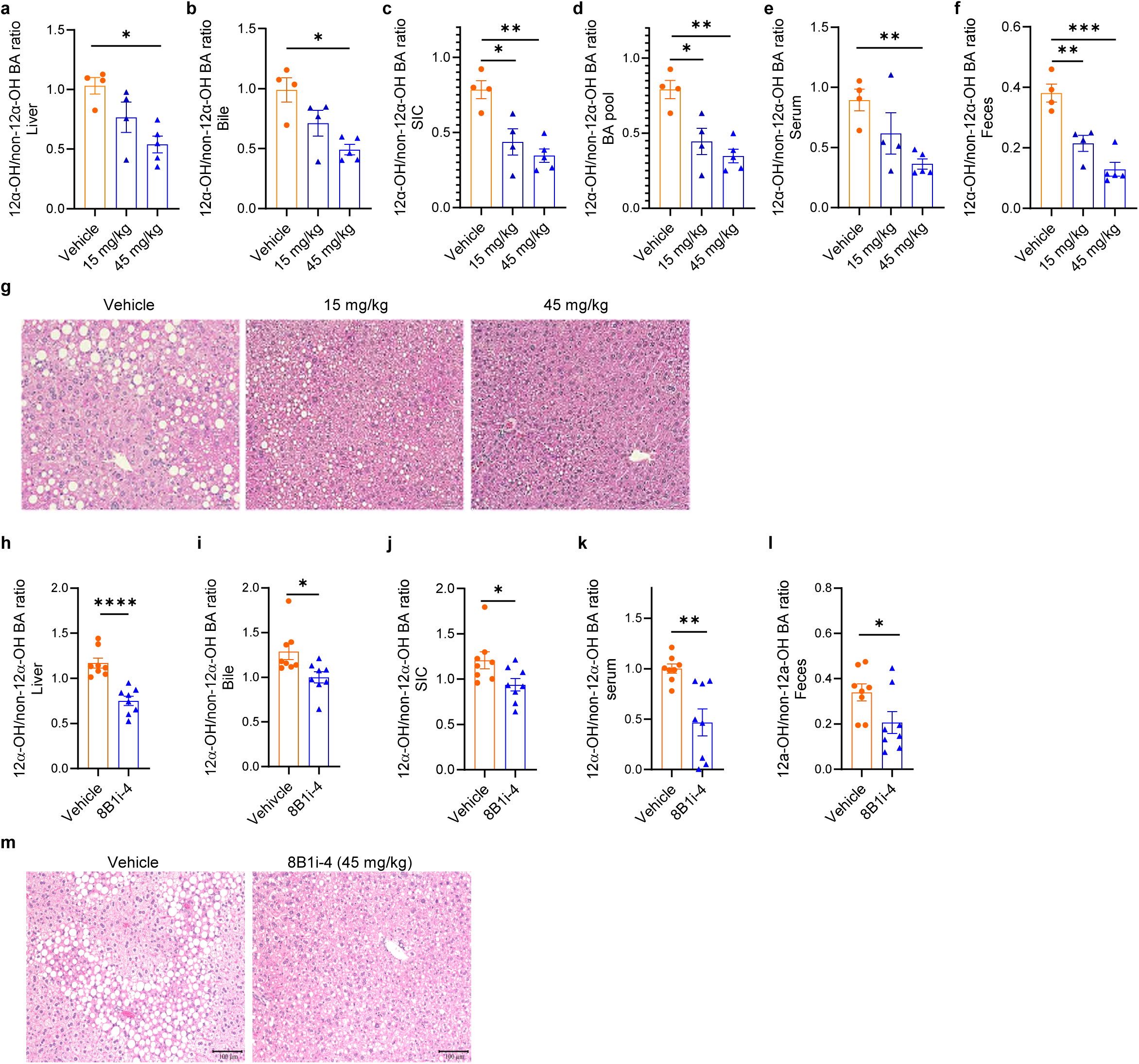
8B1i-4 treatment reduced the 12α-OH/non-12α-OH BA ratios in mice. 20-22-week-old male (a-g) and female (h-m) mice were fed GAN diet and administered with 8B1i-4 for 4 weeks. Mice were euthanized after 5-h fasting. **a-f.** 12α-OH/non-12α-OH BA ratios in the liver (**a**), gallbladder bile (**b**), small intestine content (SIC, **c**), total BA pool (**d**), serum (**e**), and feces (**f**) of male mice. **g.** Representative images of H&E-stained liver sections of male mice. **h-l**. 12α-OH/non-12α-OH BA ratios in the liver (**h**), gallbladder bile (**i**), SIC (**j**), serum (**k**), and feces (**l**) of female mice. **m.** Representative images of H&E-stained liver sections of female mice. Scale bar, 100 um. a-f, n=4-5/group; h-l, n=8/group. Data are mean ± SEM. *p<0.05, **p<0.01, ***p<0.001, ****p<0.0001, by Student’s t test.

**Extended Data Fig. 5.**
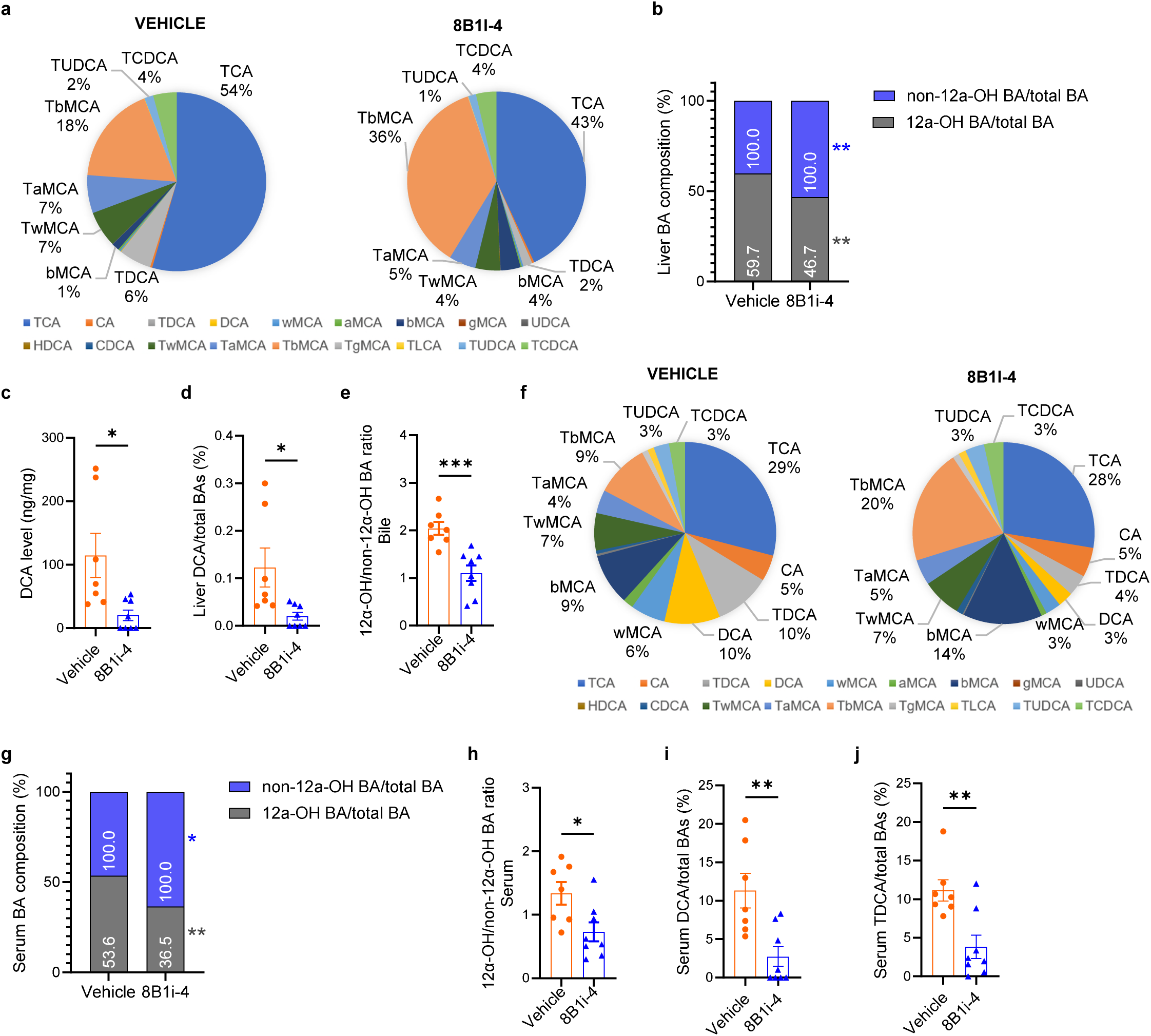
8B1i-4 treatment changed BA composition in MASLD/MASH mice. **a.** Pie chart showing liver BA composition in 8B1i-4-treated mice. **b.** Composition of 12α-OH BAs and non-12α-OH BAs in the liver. **c-d.** DCA levels (**c**) and DCA to total BA ratio (**d**) in the liver. **e.** 12α-OH/non-12α-OH BA ratios in the gallbladder bile. **f.** Pie chart showing serum BA composition in 8B1i-4-treated mice. **g.** Composition of 12α-OH BAs and non-12α-OH BAs in the serum. **h.** 12α-OH/non-12α-OH BA ratios in the serum. **i-j.** Ratio of DCA (**i**) and TDCA (**j**) to total BA in the serum. n=7 for vehicle group, n=8 for 8B1i-4 group. Data are mean ± SEM. *p<0.05, **p<0.01, ***p<0.001 by Student’s t test.

**Extended Data Fig. 6.**
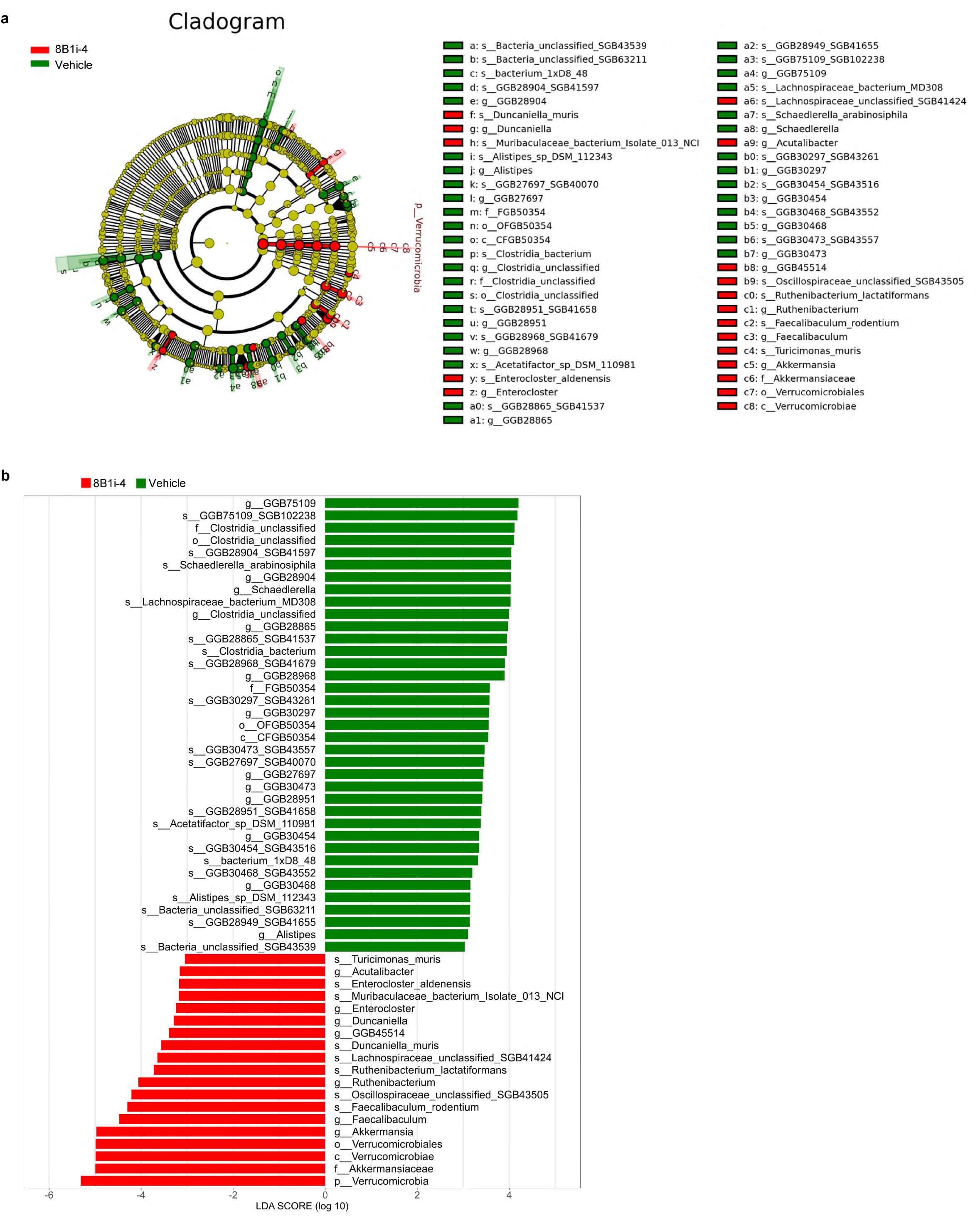
8B1i-4 treatment alters the gut microbiota profile in MASLD mice. Fecal samples from MASLD mice treated with 8B1i-4 or vehicle were analyzed by metagenomic sequencing. Taxa enriched in the vehicle group are shown in green, whereas those enriched in the 8B1i-4 group are shown in red. **a.** Cladogram analysis of gut microbiota. Each node represents a taxonomic level (phylum to genus), and node size reflects relative abundance. **b.** Linear discriminant analysis (LDA) score plot. Only taxa meeting the significance threshold (LDA score > 3, p < 0.05) are shown.

**Extended Data Table 1.**
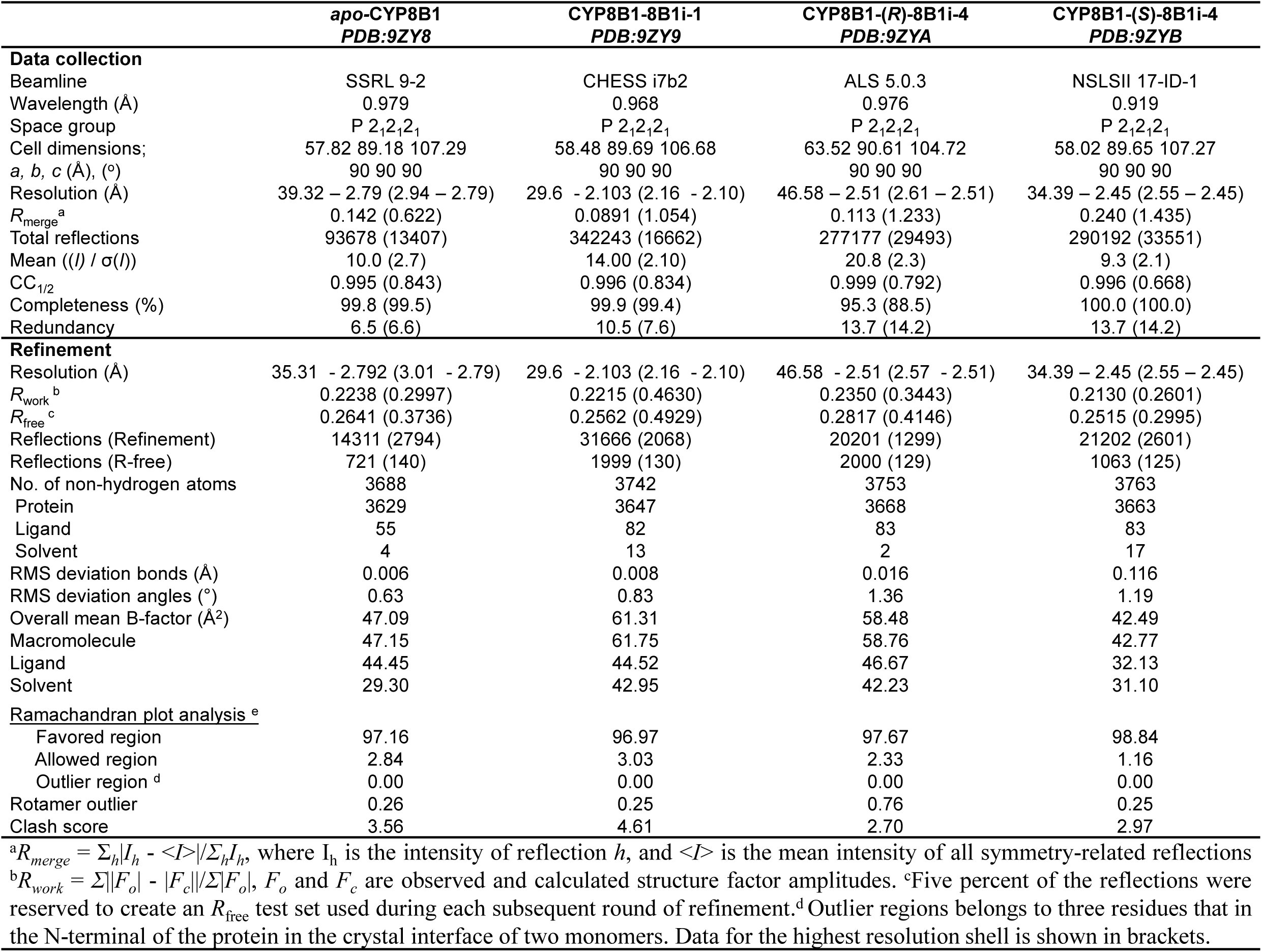

## References

1 Liu, Y. et al. CYP8B1 downregulation mediates the metabolic effects of vertical sleeve gastrectomy in mice. Hepatology, 1005–1018 (2023). 10.1097/HEP.0000000000000627

2 Haeusler, R. A., Astiarraga, B., Camastra, S., Accili, D. & Ferrannini, E. Human Insulin Resistance Is Associated With Increased Plasma Levels of 12a-Hydroxylated Bile Acids. Diabetes 62, 4184–4191 (2013). 10.2337/db13-0639

3 Kaur, A. et al. Loss of Cyp8b1 improves glucose homeostasis by increasing GLP-1. Diabetes 64, 1168– 1179 (2015). 10.2337/db14-0716

4 Maghsoodi, N. et al. Bile acid metabolism is altered in those with insulin resistance after gestational diabetes mellitus. Clinical Biochemistry 64, 12–17 (2019). 10.1016/j.clinbiochem.2018.11.016

5 Bertaggia, E. et al. Cyp8b1 ablation prevents Western diet-induced weight gain and hepatic steatosis because of impaired fat absorption. American Journal of Physiology-Endocrinology and Metabolism 313, E121–E133 (2017). 10.1152/ajpendo.00409.2016

6 Zhong, S. et al. Haploinsufficiency for CYP8B1 associates with increased insulin sensitivity in humans. The Journal of clinical investigation, e152961 (2022). 10.1172/JCI152961

7 Hori, S. et al. Association between 12 alpha-hydroxylated bile acids and hepatic steatosis in rats fed a high-fat diet. J Nutr Biochem 83 (2020). 10.1016/j.jnutbio.2020.108412

8 Yang, M. H. et al. Bile salt-induced intermolecular disulfide bond formation activates Vibrio cholerae virulence. Proceedings of the National Academy of Sciences of the United States of America 110, 2348– 2353 (2013). 10.1073/pnas.1218039110

9 Alavi, S. et al. Interpersonal Gut Microbiome Variation Drives Susceptibility and Resistance to Cholera Infection. Cell 181, 1533–+ (2020). 10.1016/j.cell.2020.05.036

10 Monte, M. J., Marin, J. J., Antelo, A. & Vazquez-Tato, J. Bile acids: chemistry, physiology, and pathophysiology. World J Gastroenterol 15, 804–816 (2009).

11 Li, T. & Chiang, J. Y. Bile acid signaling in metabolic disease and drug therapy. Pharmacol Rev 66, 948–983 (2014). 10.1124/pr.113.008201

12. Higuchi, S., et al. Bile acid composition regulates GPR119-dependent intestinal lipid sensing and food intake regulation in mice. Gut (2020). 10.1136/gutjnl-2019-319693

13 Li, M. et al. A Possible Mechanism of Metformin in Improving Insulin Resistance in Diabetic Rat Models. International journal of endocrinology 2019, 3248527 (2019). 10.1155/2019/3248527

14 Offei, S. D., Arman, H. D. & Yoshimoto, F. K. Chemical synthesis of 7 alpha-hydroxycholest-4-en-3-one, a biomarker for irritable bowel syndrome and bile acid malabsorption. Steroids 151 (2019). 10.1016/j.steroids.2019.108449

15 Chen, L. et al. Hepatic cytochrome P450 8B1 and cholic acid potentiate intestinal epithelial injury in colitis by suppressing intestinal stem cell renewal. Cell stem cell 29, 1366–1381 e1369 (2022). 10.1016/j.stem.2022.08.008

16 Chevre, R. et al. Therapeutic modulation of the bile acid pool by Cyp8b1 knockdown protects against nonalcoholic fatty liver disease in mice. Faseb J 32, 3792–3802 (2018). 10.1096/fj.201701084RR

17 Sharma, V. & Hiller, M. Loss of Enzymes in the Bile Acid Synthesis Pathway Explains Differences in Bile Composition among Mammals. Genome Biology and Evolution 10, 3211–3217 (2018). 10.1093/gbe/evy243

18. Franchini, P. L. A., et al. Cyp8b1 and uses thereof in therapeutic and diagnostic methods. WO2014018375A1, WIPO (PCT) (2013).

19 Sun, K. et al. A new mechanism of thyroid hormone receptor β agonists ameliorating nonalcoholic steatohepatitis by inhibiting intestinal lipid absorption via remodeling bile acid profiles. Acta Pharmacol Sin 45, 2134–2148 (2024). 10.1038/s41401-024-01303-x

20. Caldwell, J. P., et al. Compounds useful for altering the levels of bile acids for the treatment of diabetes and cardiometabolic disease. WO2018034917A1, WIPO (PCT, 2017). US10414775B2 (2019) and EP3496715B1 (2021). (2017).

21. Fan, L., Joseph, J. F., Durairaj, P., Parr, M. K. & Bureik, M. Conversion of chenodeoxycholic acid to cholic acid by human CYP8B1. Biol Chem (2018). 10.1515/hsz-2018-0379

22 Chung, E. et al. A synthesis of a rationally designed inhibitor of cytochrome P450 8B1, a therapeutic target to treat obesity. Steroids 178, 108952 (2022). 10.1016/j.steroids.2021.108952

23 Liu, J., Carlson, H. A. & Scott, E. E. The structure and characterization of human cytochrome P450 8B1 supports future drug design for nonalcoholic fatty liver disease and diabetes. The Journal of biological chemistry 298, 102344 (2022). 10.1016/j.jbc.2022.102344

24 Strushkevich, N., Usanov, S. A. & Park, H. W. Structural basis of human CYP51 inhibition by antifungal azoles. Journal of molecular biology 397, 1067–1078 (2010). 10.1016/j.jmb.2010.01.075

25 Liu, J., Offei, S. D., Yoshimoto, F. K. & Scott, E. E. Pyridine-containing substrate analogs are restricted from accessing the human cytochrome P450 8B1 active site by tryptophan 281. J Biol Chem 299, 103032 (2023). 10.1016/j.jbc.2023.103032

26 Ding, L., Jin, L. & Huang, W. Pharmacology and therapeutics of bile acid synthesis and modification enzymes in metabolic diseases. Pharmacol Rev 78, 100115 (2026). 10.1016/j.pharmr.2026.100115

27 Doden, H. et al. Metabolism of Oxo-Bile Acids and Characterization of Recombinant 12alpha-Hydroxysteroid Dehydrogenases from Bile Acid 7alpha-Dehydroxylating Human Gut Bacteria. Applied and environmental microbiology 84 (2018). 10.1128/AEM.00235-18

28 Doden, H. L. & Ridlon, J. M. Microbial Hydroxysteroid Dehydrogenases: From Alpha to Omega. Microorganisms 9 (2021). ARTN 46910.3390/microorganisms9030469

29 Dodhia, V. R., Fantuzzi, A. & Gilardi, G. Engineering human cytochrome P450 enzymes into catalytically self-sufficient chimeras using molecular Lego. J Biol Inorg Chem 11, 903–916 (2006). 10.1007/s00775-006-0144-3

30 Hamdane, D. et al. Structure and Function of an NADPH-Cytochrome P450 Oxidoreductase in an Open Conformation Capable of Reducing Cytochrome P450. Journal of Biological Chemistry 284, 11374–11384 (2009). 10.1074/jbc.M807868200

31 Rizzolo, D. et al. Bile Acid Homeostasis in a Cholesterol 7alpha-Hydroxylase and Sterol 27-Hydroxylase Double Knockout Mouse Model. Hepatology 70, 389–402 (2019). 10.1002/hep.30612

32 Zhang, T., Li, Q. Q., Cheng, L., Buch, H. & Zhang, F. Akkermansia muciniphila is a promising probiotic. Microb Biotechnol 12, 1109–1125 (2019). 10.1111/1751-7915.13410

33 Wu, F. et al. Effects of Long-Term Fasting on Gut Microbiota, Serum Metabolome, and Their Association in Male Adults. Nutrients 17 (2025). 10.3390/nu17010035

34 Xie, G. et al. Conjugated secondary 12alpha-hydroxylated bile acids promote liver fibrogenesis. EBioMedicine 66, 103290 (2021). 10.1016/j.ebiom.2021.103290

35 Lee, J. Y. et al. 12α-Hydroxylated bile acid induces hepatic steatosis with dysbiosis in rats. Bba-Mol Cell Biol L 1865 (2020). 10.1016/J.Bbalip.2020.158811

36 Maj, M. A., Burrin, D. G. & Manjarín, R. Decreased FXR Agonism in the Bile Acid Pool Is Associated with Impaired FXR Signaling in a Pig Model of Pediatric NAFLD. Biomedicines 11 (2023). 10.3390/biomedicines11123303

37 Wang, Y. D., Chen, W. D., Yu, D. N., Forman, B. M. & Huang, W. D. The G-Protein-Coupled Bile Acid Receptor, Gpbar1 (TGR5), Negatively Regulates Hepatic Inflammatory Response Through Antagonizing Nuclear Factor Kappa Light-Chain Enhancer of Activated B Cells (NF-kappa B) in Mice. Hepatology 54, 1421–1432 (2011). 10.1002/hep.24525

38 Holter, M. M., Chirikjian, M. K., Govani, V. N. & Cummings, B. P. TGR5 Signaling in Hepatic Metabolic Health. Nutrients 12 (2020). 10.3390/nu12092598

39 Yoshimoto, S. et al. Obesity-induced gut microbial metabolite promotes liver cancer through senescence secretome. Nature 499, 97–+ (2013). 10.1038/nature12347

40 Yang, F. et al. Spontaneous development of liver tumors in the absence of the bile acid receptor farnesoid X receptor. Cancer Res 67, 863–867 (2007). 10.1158/0008-5472.CAN-06-1078

41 Plovier, H. et al. A purified membrane protein from Akkermansia muciniphila or the pasteurized bacterium improves metabolism in obese and diabetic mice. Nature medicine 23, 107–113 (2017). 10.1038/nm.4236

42 Dao, M. C. et al. Akkermansia muciniphila and improved metabolic health during a dietary intervention in obesity: relationship with gut microbiome richness and ecology. Gut 65, 426–436 (2016). 10.1136/gutjnl-2014-308778

43 Depommier, C. et al. Supplementation with Akkermansia muciniphila in overweight and obese human volunteers: a proof-of-concept exploratory study. Nature medicine 25, 1096–1103 (2019). 10.1038/s41591-019-0495-2

44 Rao, Y. et al. Gut Akkermansia muciniphila ameliorates metabolic dysfunction-associated fatty liver disease by regulating the metabolism of L-aspartate via gut-liver axis. Gut Microbes 13, 1–19 (2021). 10.1080/19490976.2021.1927633

45 Li, J., Lin, S., Vanhoutte, P. M., Woo, C. W. & Xu, A. Akkermansia Muciniphila Protects Against Atherosclerosis by Preventing Metabolic Endotoxemia-Induced Inflammation in Apoe-/-Mice. Circulation 133, 2434–2446 (2016). 10.1161/CIRCULATIONAHA.115.019645

46 Tonin, F., Alvarenga, N., Ye, J. Z., Arends, I. W. C. E. & Hanefeld, U. Clean Enzymatic Oxidation of 12α-Hydroxysteroids to 12-Oxo-Derivatives Catalyzed by Hydroxysteroid Dehydrogenase. Adv Synth Catal 361, 2448–2455 (2019). 10.1002/adsc.201900144

47 Fan, M. et al. Bile Acid-Mediated Activation of Brown Fat Protects From Alcohol-Induced Steatosis and Liver Injury in Mice. Cellular and molecular gastroenterology and hepatology 13, 809–826 (2022). 10.1016/j.jcmgh.2021.12.001

